# A network of CD163^+^ macrophages monitors enhanced permeability at the blood-dorsal root ganglion barrier

**DOI:** 10.1101/2023.03.27.534318

**Authors:** Harald Lund, Matthew Hunt, Zerina Kurtovic, Katalin Sandor, Noah Fereydouni, Anais Julien, Christian Göritz, Jinming Han, Keying Zhu, Robert A. Harris, Jon Lampa, Lisbet Haglund, Tony L. Yaksh, Camilla I. Svensson

## Abstract

In dorsal root ganglia (DRG), macrophages reside in close proximity to sensory neurons, and their functions have largely been explored in the context of pain, nerve injury and repair. In this study, however, we discovered that the majority of macrophages in DRGs are in direct contact with the vasculature where they constantly monitor the circulation, efficiently phagocytosing proteins and macromolecules from the blood. Characterization of the DRG endothelium revealed a specialized vascular network spanning the arteriovenous axis, which gradually transformed from a barrier type endothelium in arteries to a highly permeable endothelium in veins. Macrophage phagocytosis spatially aligned with peak endothelial permeability and we identified caveolar transcytosis as a mechanism regulating endothelial permeability. Profiling of the DRG immune landscape revealed two subsets of perivascular macrophages with distinct transcriptome, turnover and function. CD163 expressing macrophages self-maintained locally, specifically participated in vasculature monitoring, displayed distinct responses during peripheral inflammation and were conserved in mouse and Man. Our work provides a molecular explanation for the permeability of the blood-DRG barrier and identifies an unappreciated role of macrophages as integral components of the DRG-neurovascular unit.

## Introduction

Tissue-resident macrophages are multi-functional and highly plastic immune cells that are found in every organ of the body. A core function of macrophages shared across most organs is to act as tissue sentinels and scavengers, by phagocytosing cellular debris and orchestrating tissue repair. In addition, macrophages participate in more complex, tissue-specific processes, as diverse as wiring of neural networks, production of blood vessels and bone as well as turnover of dying cells or essential proteins in the liver, spleen and lung (Park et al., 2022; Mass et al., 2023). It is now understood that such tissue-specialized functions are the result of distinct molecular programs induced by microenvironmental cues (cell-contacts and secreted molecules) in their tissues of residence (Gosselin et al., 2014; Lavin et al., 2014; Bonnardel et al., 2019). Another factor contributing to macrophage variability in different organs relates to their ontogeny. During development, organs are colonized by embryonic macrophages derived from precursors in the yolk sac or fetal liver that can self-sustain throughout life (Ginhoux et al., 2010; Schulz et al., 2012; Hoeffel et al., 2015; Perdiguero et al., 2015). By contrast, there are subpopulations of macrophages identified in most tissues which require constant replenishment from bone-marrow derived monocytes (Molawi et al., 2014; Shaw et al., 2018; Dick et al., 2022).

Macrophages in the CNS are well-studied, where parenchymal microglia and border-associated macrophages, residing in perivascular, meningeal and choroid plexus niches, are recognized (Mildenberger et al., 2022). In the peripheral sensory nervous system, however, aspects such as transcriptional heterogeneity, ontogeny, microenvironmental regulation and homeostatic function are only starting to be explored (Kolter et al., 2020; Wang et al., 2020a; Ydens et al., 2020). Macrophages associate with the entire length of sensory neurons, from the distal peripheral nerve endings to the proximal processes entering the spinal column. Macrophages are also prominent in dorsal root ganglia (DRG) (Zigmond and Echevarria, 2019), which are segmentally organized collections of sensory neuron cell bodies located alongside the spinal cord. Transcriptional differences between macrophages residing in these different locations are evident, indicating that they are a product of their endoneurial microenvironment (Wang et al., 2020a). DRG macrophages increase in number during a range of neuropathic conditions (Peng et al., 2016; Zhang et al., 2016; Vlist et al., 2022), which may promote the development of pain (Yu et al., 2020; Raoof et al., 2021) or help resolve it (Singh et al., 2022; Vlist et al., 2022). However, the homeostatic functions of macrophages in the DRG have remained largely overlooked. Moreover, heterogeneity within the macrophage pool, which is recognized in virtually every organ investigated, has not been systematically examined in the DRG.

The axons of sensory neurons are protected from circulating toxic molecules and pathogens by the blood-nerve barrier (BNB), which shares functional and morphological features with the blood-brain barrier (BBB), including low levels of transcytosis, high expression of tight junction proteins and lack of endothelial fenestrae (Ubogu, 2020). The neuronal cell bodies in the DRG, however, are not only supplied by a denser vascular bed than the axons in the peripheral nerve (Jimenez-Andrade et al., 2008), the endothelial cells within DRGs also have significantly higher permeability (Olsson, 1968; Jacobs et al., 1976; Arvidson, 1979; Kobayashi and Yoshizawa, 2002). Despite the long-standing appreciation for these properties, an understanding of the mechanisms regulating endothelial permeability at the blood-DRG barrier remain lacking. Furthermore, the role of macrophages in this context has remained unexplored. Breakdown of endothelial barrier integrity occurs and contributes to disease progression in a range of conditions affecting the BBB (Profaci et al., 2020) and the BNB (Richner et al., 2019; Ubogu, 2020). The elevated permeability observed in DRG endotheliu makes the DRG susceptible to neurotoxic molecules, autoantibodies and infectious agents, and is likely a major cause of sensory ganglionopathies, conditions leading to sensory loss or painful manifestations (Amato and Ropper, 2020). Characterizing the cellular components of the vascular niche and understanding how endothelial barrier integrity is maintained in the DRG therefore remain important areas of research.

In this study we describe a dense network of perivascular macrophages residing in sensory ganglia, tasked with monitoring the permeable blood-DRG barrier. Mechanistically, we found that this process was driven by elevated caveolar transcytosis in endothelial cells coupled to a self-sustained and highly phagocytic macrophage subset. Our work identifies a novel immunovascular unit that has implications for understanding blood-nerve barrier homeostasis, disease and therapy.

## Results

### DRG macrophages monitor the vasculature

Macrophage functions are shaped by and tailored to their cellular microenvironment, known as the ‘macrophage niche’ (Guilliams et al., 2020). To characterize the macrophage niche in the DRG at a cellular level, we co-stained macrophages (Iba1^+^) and major cell types in the DRG, including neurons (Neurotrace^+^), and satellite glial cells (SGC, GS^+^) which wrap around the neuronal soma (Hanani and Spray, 2020). Given the high degree of vascularization in the DRG (Jimenez-Andrade et al., 2008), we also stained endothelial cells. This analysis revealed that macrophages were positioned in the space between SGCs and endothelial cells (Fig. 1a), often making close contact with endothelial cells (Fig. 1b). This prompted us to further explore the interaction between macrophages and endothelial cells in the DRG.

**Figure 1:**
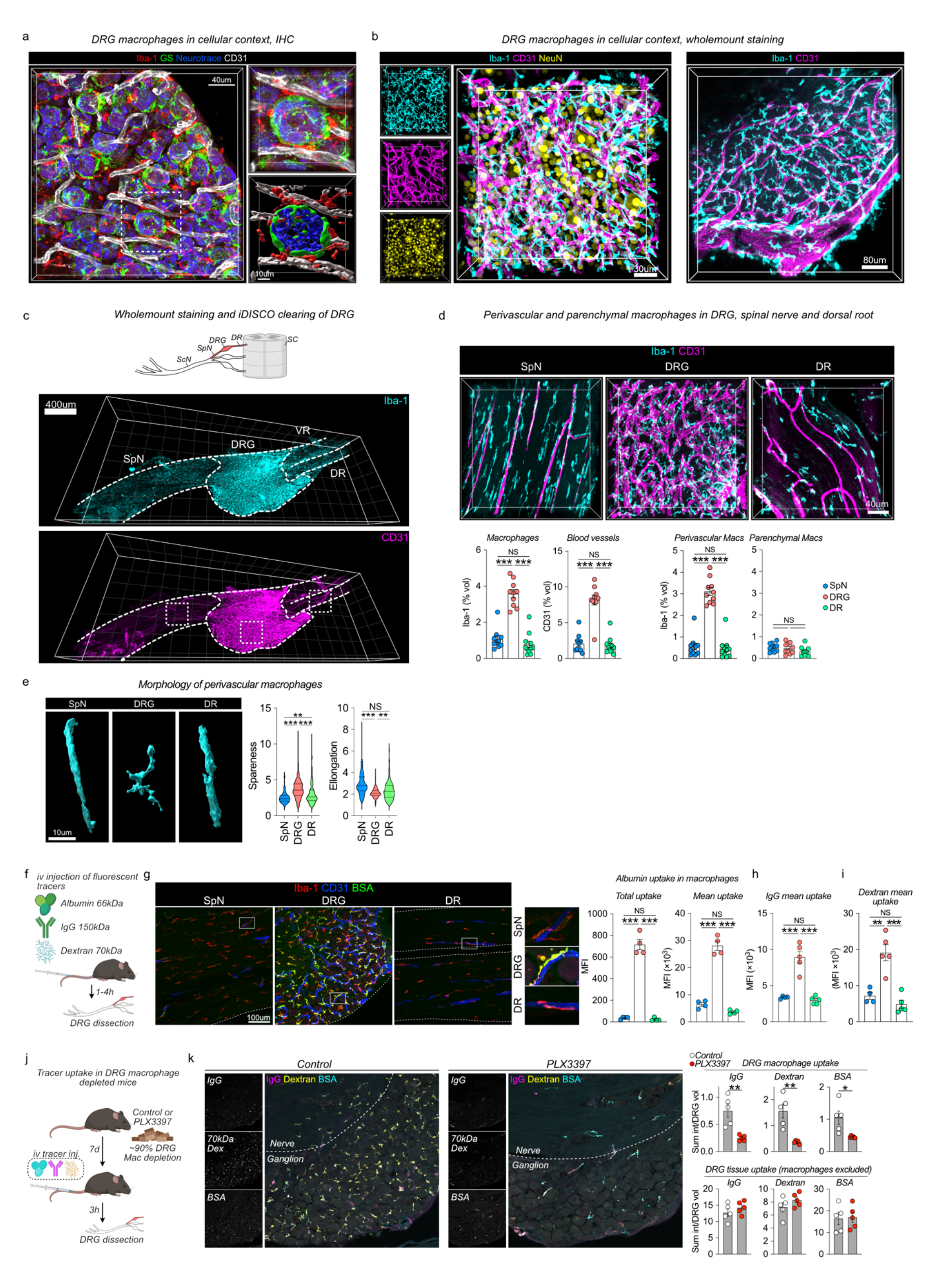
**DRG macrophages interact closely with the vasculatur** (a) Representative immunostaining of neurons, SGCs, endothelial cells and macrophages in naive DRGs. (b) 3D confocal images of DRG whole mounts to visualize neuron, macrophage and endothelial cells. (c) 3D Confocal images of iDISCO-cleared whole mounts of DRGs with attached spinal nerve (SpN) and dorsal/ventral roots (DR/VR) stained with Iba-1 and CD31. (d) Confocal Z-stacks of boxed areas in (c). Quantifications of macrophage and vascular density performed in indicated regions. n=10, 10, 11 mice/group. Tukey’s multiple comparisons test. (e) Morphology of perivascular macrophages as measured by their spareness (high value indicates spider-like shape) and elongation (high value indicates cigar-shape). Tukey’s multiple comparisons test. Experiment schematic of i.v injection of fluorescent tracers. (f) BSA-A647 uptake in Iba-1^+^ macrophages 1h after i.v injection. N=4mice/group. The experiment was performed twice. Tukey’s multiple comparisons test. (g) Uptake of goat IgG-A488 (4mg/kg) in Iba-1^+^ macrophages 4h after i.v injection. N=4mice/group. The experiment was performed twice. Tukey’s multiple comparisons test. (h) Uptake of 70kDa dextran-TMR in Iba-1^+^ macrophages 2h45min after i.v injection. The experiment was performed twice. 1-way ANOVA and Tukey post-hoc test. (i) Experiment schematic of i.v tracer uptake in macrophage depleted mice using the CSF1R antagonist PLX3397 (290ppm in chow). Tukey’s multiple comparisons test. (j) Uptake of BSA-A647 (4 mg/kg), 70kDa dextran-TMR (10mg/kg) and goat anti rabbit IgG-A488 (3mg/kg) in Iba-1^+^ macrophages or in DRG parenchyma (excluding macrophages) 2h45min after i.v injection. The experiment was performed once. Student’s unpaired t-test.

We next stained Iba-1^+^ macrophages and CD31^+^ endothelial cells in DRG wholemounts, followed by iDISCO tissue clearing and 3D-confocal imaging (Fig. 1c). DRGs were dissected with the spinal nerve (SpN) and dorsal root (DR) attached, which allowed comparisons of macrophages across these three tissue regions. Quantification of macrophage volume showed a significant increase in the DRG compared to adjacently located SpN and DR (Fig. 1d). This finding was confirmed using flow cytometry of enzymatically digested tissues, showing that the DRG contained a higher number of CX3CR1^+^CD64^+^ macrophages than did the sciatic nerve (ScN) per weight of tissue (Supplementary Fig. 1a). Comparison of the level of vascularization similarly demonstrated a 4-fold increase in the DRG compared to the SpN and DR (Fig. 1d), which supports previous findings (Jimenez-Andrade et al., 2008). We then investigated the spatial relationship between endothelial cells and macrophages, observing that macrophages that were in direct contact with the abluminal side of endothelial cells (perivascular) were approximately 6-fold more prevalent in the DRG than in the DR or SpN (Fig. 1d). Macrophages that did not make contact with the vasculature (parenchymal) were similar in volume across the three tissue regions (Fig. 1d). The morphology of perivascular DRG macrophages was also distinct, coiling around vessels and displaying a more tortuous shape than did their SpN and DR counterparts. SpN and DR perivascular macrophages had a more elongated shape, extending along blood vessels parallel to the axons (Fig. 1e). Our data thus far demonstrate a local increase in perivascular macrophages specifically in the DRG.

Given the well-described capacity of low and high molecular weight compounds to permeate the blood-DRG barrier (Olsson, 1968; Jacobs et al., 1976; Arvidson, 1979; Kobayashi and Yoshizawa, 2002), we next addressed whether macrophages had the capacity to phagocytose circulating molecules. To that end we intravenously injected mice with either fluorescently labeled albumin (BSA, 66kDa) or IgG (150 kDa), the two most abundant proteins in plasma. Sacrificing animals within 1 hour (BSA) or 4 hours (IgG) revealed that both proteins readily accumulated inside DRG macrophages and quantification across sensory nerve regions demonstrated a 4-to-7 fold (BSA) or 2-fold (IgG) higher uptake in DRG macrophages compared to in SpN or DR macrophages (Fig. 1f-h). To substantiate these findings we next injected 70kDa dextran, a branched glucan used clinically as a plasma substitute that displays minimal extravasation across the healthy BBB(Armulik et al., 2010). Similarly to our IgG and BSA injections, dextran uptake was elevated in DRG macrophages compared to in SpN and DR macrophages (Fig. 1i). Next we addressed whether DRG macrophages were actively monitoring the circulation or only passively phagocytosing material leaking through the blood-DRG barrier. Administration of the CSF1R-antagonist PLX3397 in chow resulted in 90% depletion of DRG macrophages within 7 days (Supplementary Fig. 1b, Fig. 1j) and as expected, resulted in abrogation of vascular monitoring of IgG, dextran and BSA (Fig. 1k). However, we did not observed increased tracer leakage into the DRG parenchyma (Fig. 1k), indicating that macrophages were actively involved in the monitoring process. These results demonstrate that the majority of DRG macrophages interact with the vasculature and monitor the circulation for both endogenous and exogenous molecules.

### The DRG vasculature displays both barrier and permeable properties and has a conserved arteriovenous distribution

While differences in vascular permeability between the blood-nerve and blood-ganglion barrier are recognized (Reinhold and Rittner, 2020), an in-depth molecular understanding of this phenomenon is lacking. We hypothesized that increased vascular permeability could at least partly explain the level of circulating protein uptake in DRG macrophages, and we next sought to better characterize DRG endothelial cells. Guided by a previous report (Munji et al., 2019), we first analyzed whether DRG endothelial cells expressed markers specific to the BBB or peripheral endothelium. This revealed that DRG endothelial cells expressed high levels of the glucose transporter *Slc2a1* as well as the amino acid transporter *Slc7a5*, both which are specific to the BBB (Munji et al., 2019; Kalucka et al., 2020). DRG endothelial cells also expressed *Gpihbp1*, involved in lipid metabolism, and the prostaglandin transporter *Slco2a1*, these genes normally being expressed in the kidney, liver and lung endothelium but are absent from the BBB (Supplementary Fig. 2a). These results indicated that DRG blood vessels expressed both peripheral and CNS-specific markers, and further suggested transcriptomic heterogeneity across the DRG endothelial population.

Progressive transcriptomic changes along the arteriovenous axis, a process defined as ‘zonation’, has been recognized in several endothelial beds using scRNA-seq (Vanlandewijc et al., 2018; Kalucka et al., 2020). To address transcriptomic zonation in the DRG vasculature, we re-clustered a publicly available mouse scRNA-seq dataset (Avraham et al., 2020) which included 432 DRG endothelial cells. Three clusters of endothelial cells were identified (Fig. 2b), which we annotated as *artery, capillary* and *vein*, respectively, based on expression of well-established (Vanlandewijck et al., 2018; Kalucka et al., 2020; Trimm and Red-Horse, 2022) artery-(*Hey1, Bmx, Vegfc, Sema3g*) and vein-specific markers (*Nr2f2, Bgn, Vcam1, Vwf*) in the two clusters that occupied the extremes of the UMAP (Supplementary Fig. 2b). Differential gene expression further revealed that the artery cluster was characterized by high expression of *Cldn5, Slc2a1* and *Mfsd2a*, which are all highly enriched in brain endothelial cells (Supplementary Fig. 2c-d). *Cldn5* encodes a tight junction protein which maintains BBB integrity (Greene et al., 2019) and *Mfsd2a* is a transporter of essential fatty acids required for proper brain development and function (Nguyen et al., 2014). The vein cluster displayed high expression of *Plvap, Aqp1, Gpihbp1* and *Lrg1*, all being enriched in peripheral endothelial beds (Supplementary Fig. 2c-d). *Plvap* encodes a protein restricted to endothelial fenestrae, transendothelial channels and caveolar vesicles; structures involved in microvascular permeability (Guo et al., 2016). All these markers, including *Plvap* and *Cldn5*, displayed a zonated expression profile, peaking in either arteries or veins and gradually decreasing or increasing along the arteriovenous axis (Fig. 2c).

**Figure 2:**
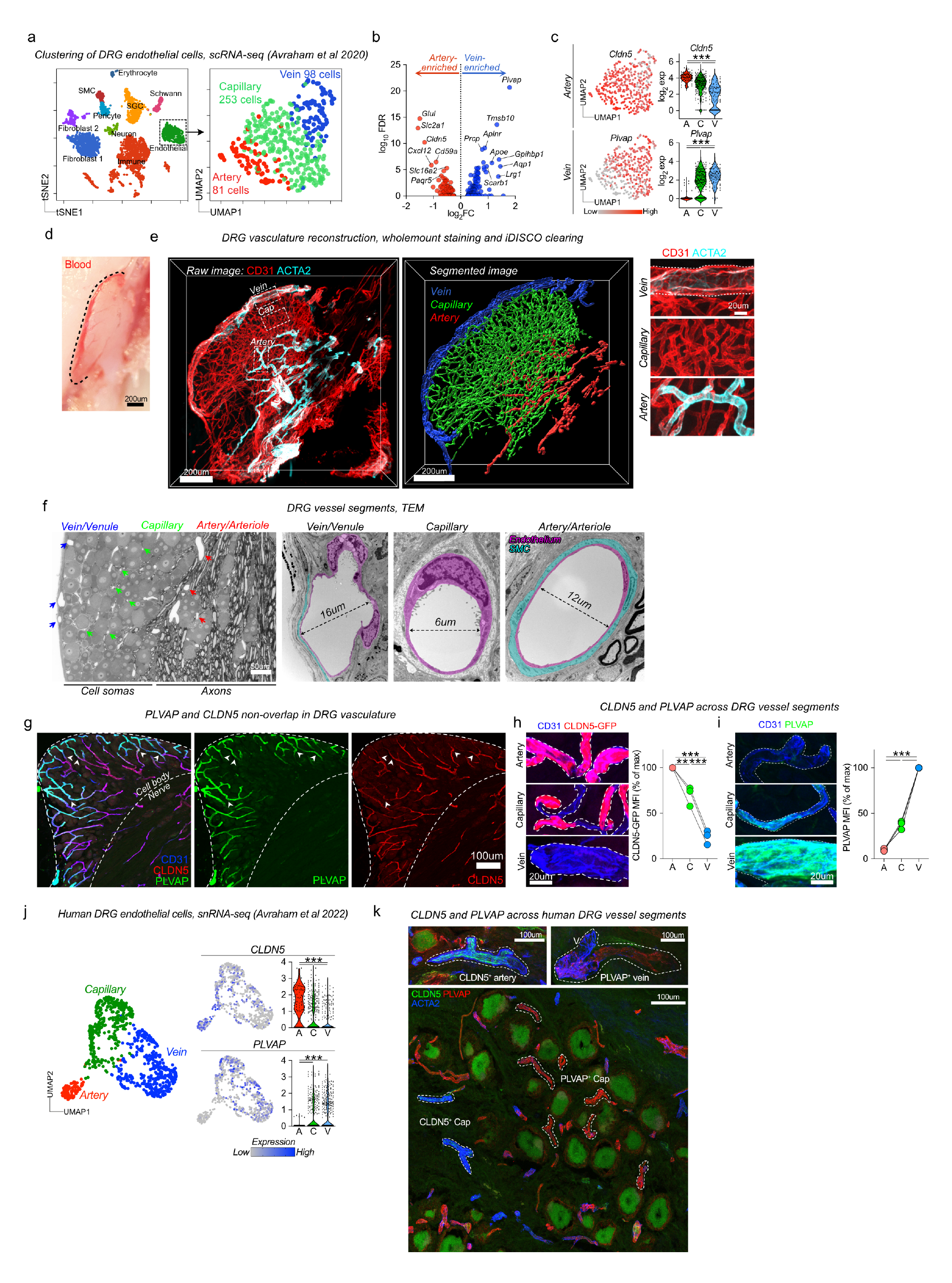
**DRG vasculature has dual identity** (a) Analysis of 432 endothelial cells from Avraham et al displaying clusters with artery, capillary and vein identity. (b) Differential gene expression between artery and vein clusters using Venice algorithm. (c) *Cldn5* (artery) and *Plvap* (vein) expression across clusters. (d) Photomicrograph of undissected L5 DRG from unperfused mouse illustrating blood-filled vasculature. (e)Whole mount imaging and iDISCO tissue-clearing of CD31 and ACTA2 on lumbar DRG. 3D-reconstruction and vessel segment identification using Imaris. (f) Identification of vessel segments in ultrathin DRG sections by TEM, based on their anatomical localization. (g) CLDN5 and PLVAP expression in CD31^+^ DRG endothelial cells, displaying minimal overlap. Arrows indicate PLVAP/CLDN5 breakpoints. (h) Mean CLDN5 expression in DRG vessel segments, normalized to % of max. n=3 mice. Tukey’s multiple comparisons test. (i) Mean PLVAP expression in DRG vessel segments, normalized to % of max. n=3 mice. Tukey’s multiple comparisons test. (j) Clustering of 777 endothelial cells from human DRGs from 5 donors (Avraham et al 2022) and the expression of *CLDN5* and *PLVAP* across artery, capillary and vein clusters. (k) Immunostaining of CLDN5 and PLVAP in human DRG sections. A= artery. V=vein. Cap = Capillary

We next wanted to validate these markers at the protein level. In order to anatomically identify the three vessel segments we stained CD31 and smooth muscle actin (SMA/ACTA2) in DRG wholemounts and used 3D-confocal imaging to reconstruct the DRG vasculature. This analysis revealed a conserved anatomical distribution, where arteries entered the neuron-rich region from the nerve fiber, giving rise to a capillary bed that was subsequently collected into veins on the DRG surface (Fig. 2d-e). Transmission electron microscopy (TEM) further validated the anatomical location of these vessel segments (Fig. 2f). We explored the expression of the two top artery-and vein-markers, *Cldn5* and *Plvap*, at the protein level. Co-staining in DRG sections revealed that both markers were restricted to endothelial cells, but with minimal overlap, PLVAP being expressed in superficially located vessels and CLDN5 in those closer to the nerve fiber (Fig. 2g). Using *Cldn5*^GFP/+^ mice we could confirm high expression of CLDN5 in ACTA2^+^ arteries, intermediate expression in capillaries, and complete absence in large veins on the DRG surface (Fig. 2h, Supplementary Fig. 2e, Supplementary Fig. 2g). This pattern was completely reversed for PLVAP, which displayed the highest expression in large veins, intermediate expression in capillaries whereas ACTA2^+^ arteries were completely devoid of PLVAP expression (Fig. 2i, Supplementary Fig. 2f). The presence of CLDN5^−^PLVAP^+^ capillaries entering the endoneurium appeared unique to the DRG, as it was not observed in the SpN or ScN, where CLDN5^−^PLVAP^+^ vessels were restricted to the epineurium (Supplementary Fig. 2h). Our results reveal that the DRG arteriovenous tree has a predictable anatomical localization and is distinguished by a gradual phenotypic shift characterized by loss of barrier properties and gain of permeability properties.

### Arteriovenous zonation is conserved in human DRG vasculature

Human DRGs have higher *in vivo* blood perfusion rates than the SpN as measured by fMRI (Godel et al., 2016), suggesting that DRG endothelial permeability is a feature shared between mouse and Man. However, the vasculature has not previously been studied in detail in human DRGs. We thus sought to address whether human DRGs displayed a similar profile as in mouse. We first analyzed a recently published snRNA-seq dataset of human DRGs (Avraham et al., 2022), which included one cluster of 777 *PECAM1*^+^ (encoding CD31) endothelial cells. Subclustering of endothelial cells revealed three distinct clusters that could be assigned vein, capillary and artery annotations, based on expression of artery (*GJA5*, *SEMA3G*, *VEGFC*) and vein (*NR2F2*, *KCNIP4*, *IL1R1*) enriched markers identified in other organs (Supplementary Fig. 2i) (Chen et al., 2022; Trimm and Red-Horse, 2022; Yang et al., 2022). Exploring the top barrier and permeability markers identified in mouse DRG endothelial cells, *CLDN5* and *PLVAP*, revealed that both genes were zonated and similarly enriched in arteries and veins, respectively (Fig. 2j). Using DRG tissues collected from human organ donors of varying sex and age (Supplementary Table 1), we validated CLDN5 expression in ACTA2^+^ arteries and arterioles and its absence from veins and most capillaries supplying the neuronal soma-rich region (Fig. 2k). Consistent with the snRNA-seq data, PLVAP was absent from arteries, but stained most capillaries supplying the neuronal soma-rich areas, as well as large veins in the capsule (Fig. 2k). This indicates that the human DRG vasculature is also characterized by a dual identity with barrier-type arteries and highly permeable veins.

### Macrophage monitoring of DRG vasculature is arteriovenously zonated and requires caveolar transcytosis

The zonated proteogenomic profile of the DRG vasculature next led us to address whether endothelial permeability was variable along the arteriovenous axis. Using DRGQuant, a machine-learning based algorithm that we recently developed to analyze DRG macrophages in tissue sections (Hunt et al., 2022) we performed spatial mapping of macrophages along the arteriovenous tree (Fig. 3a). We intravenously injected mice with a series of macromolecules of different size including dextrans, albumin and IgG, ranging from 3 kDa to 2000 kDa, and analyzed macrophage-mediated uptake and organized the data based on which vessel segment the macrophages contacted. This demonstrated a gradual increase in macrophage uptake from arteries to veins, consistently peaking in either venous capillaries (v-cap) or veins (Fig. 3a). While the mean uptake in endothelial cells was lower than in macrophages, the uptake across vessel segments mirrored that in macrophages (Supplementary Fig. 3a).

**Figure 3:**
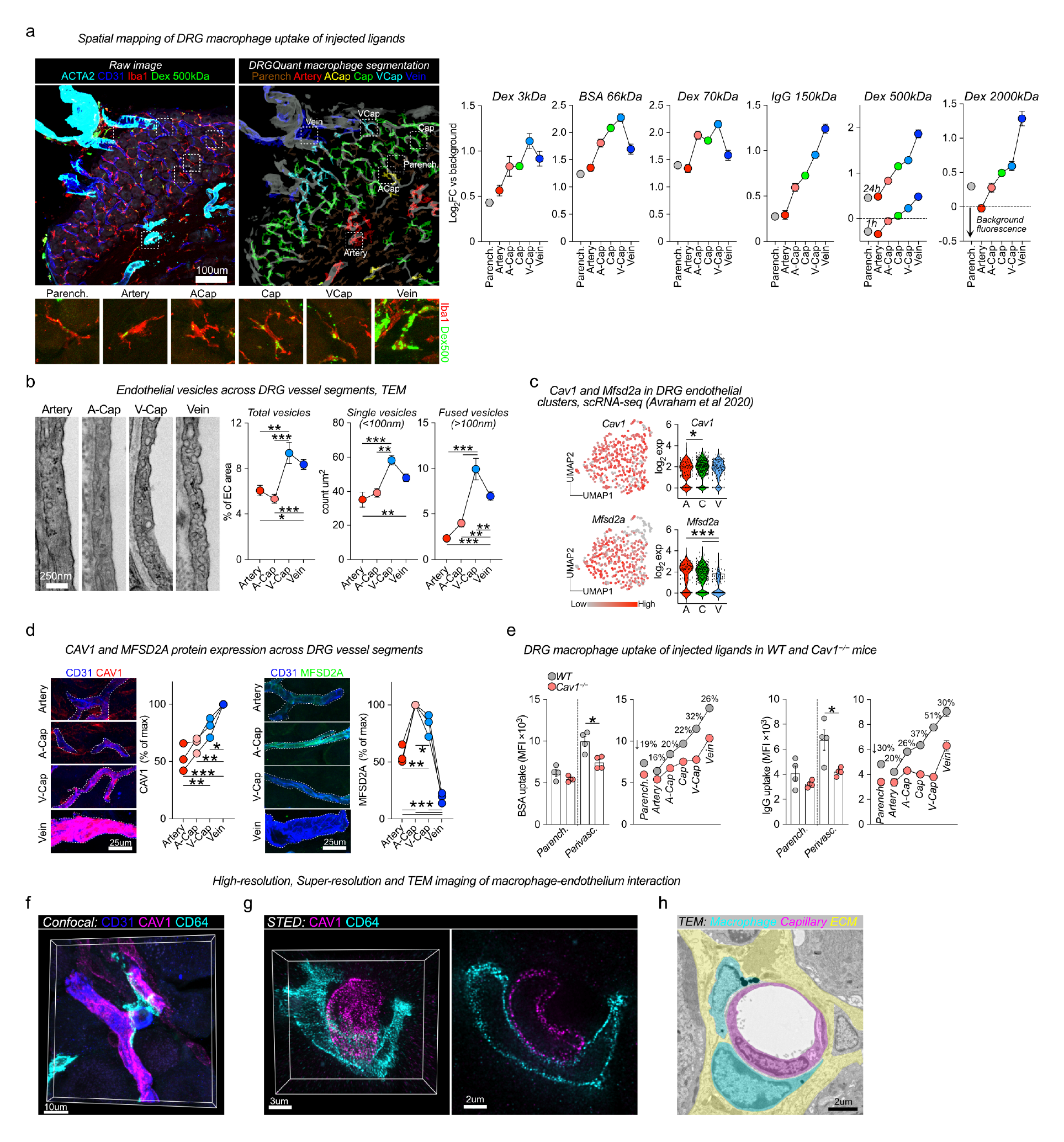
**Macrophage monitoring is arteriovenously zonated and requires caveolar vesicles** (a) Machine learning-based spatial mapping of Iba-1^+^ macrophages to DRG vessel segments and analysis of uptake of indicated i.v-injected tracers, using the DRGQuant algorithm. The following tracers, doses and circulation times were used: 3kDa dextran-TMR (25mg/kg, 1h, n=4 mice), BSA-A647 (5mg/kg, 1h, n=4 mice), 70kDa dextran TMR (25mg/kg, 1h, n=9 mice), goat anti rabbit IgG-A488 (4mg/kg, 4h, n=4 mice), 500kDa dextran-FITC (25mg/kg, 1h or 24h, n=4 mice), 2000kDa dextran-FITC (25mg/kg, 24h, n=4 mice). Values are mean of individual macrophages, normalized to tissue background. (b) Machine learning-based quantifications of endothelial vesicles (<100nm diameter) in high-resolution images of indicated DRG vessel segments. Data are mean of 50 (A), 51 (A-Cap), 42 (V-Cap) and 120 (V) images from n= 2 mice. Tukey’s multiple comparisons test. (c) Expression of *Cav1* and *Mfsd2a* mRNA in DRG vessel segments. Tukey’s multiple comparisons test. (d) Immunostaining of CAV1 and MFSD2A in DRG sections, mean intensity analyzed in indicated vessel segments. n=3 mice Tukey’s multiple comparisons test. (e) i.v injection of BSA-A488 (1mg/ml) and Goat IgG-A647 (1mg/ml) in WT and *Cav1*^−/−^ mice (n=4/group), sacrificed after 2h. Bar graphs are quantification of uptake in parenchymal and perivascular Iba-1^+^ macrophages. Line-connected graphs are quantifications of perivascular macrophages across endothelial vessel segments. Percentages indicate the reduction in macrophage-uptake between WT to *Cav1*^−/−^ mice at each vessel segment. Multiple unpaired t-test with Holm Sidak correction. (f) Confocal image of CD64^+^ macrophage with CAV1^+^ DRG blood vessel. (g) STED-captured Z-stack (left) and one Z-layer (right). (h) TEM image of macrophage-endothelial contact.

The DRG vasculature thus displayed arteriovenously zonated permeability which correlated with the presence CLDN5^−^PLVAP^+^ endothelial cells. As PLVAP is restricted to endothelial fenestrae, transendothelial channels and caveolae (Guo et al., 2016), we next sought to quantify the presence of these substructures across DRG vessel segments using TEM. The presence of fenestrael openings in the DRG vasculature have been reported (Anzil et al., 1976; Jacobs et al., 1976; Arvidson, 1979; Kobayashi and Yoshizawa, 2002), and we did identify fenestrae which were virtually restricted to v-caps (Supplementary Fig. 3b), although their numbers were limited (only 0.06% of endothelial lining). The overall scarcity of fenestrae was further confirmed by scanning electron microscopy (SEM) visualization of the inner lumen of DRG vessels (Supplementary Fig. 3c). We did not observe any transendothelial channels in the DRG vasculature (data not shown). Using higher magnification TEM images, we instead observed that small endothelial vesicles (∼100nm), likely caveolar vesicles, were ubiquitous in the DRG vasculature. Machine-learning based image quantification revealed a significantly higher presence of such vesicles in v-caps and veins compared to in arteries and arterial capillaries (a-caps) (Fig. 3b). We confirmed that these vesicles were caveolar vesicles, as they were absent in the DRG endothelium from *Cav1*^-/-^ mice (Supplementary Fig. 3d) which cannot form caveolae (Parton et al., 2020). However, fenestrae were still present in DRG v-caps from *Cav1*^-/-^ mice (Supplementary Fig. 3d). To further understand the regulation of caveolae across DRG vessel segments we quantified expression of CAV1, the main scaffolding protein required for caveolae assembly. We also explored expression of MFSD2A, an inhibitor of caveolar transcytosis (Andreone et al., 2017) which is restricted to barrier endothelium in the CNS, testis (Supplementary Fig. 3e) and retina (Wang et al., 2020b). We recorded high levels of Mfsd2a mRNA and protein in DRG endothelial cells (Supplementary Fig. 3f-g), and that mRNA expression gradually decreased from arteries to veins (Fig. 3c). Protein levels of MFSD2A were similarly zonated, reaching a peak in a-caps and then gradually decreasing to negligible levels in veins (Fig. 3d, Supplementary Fig. 3g, Supplementary Fig. 3i). *Cav1* was not zonated at the mRNA level (Fig. 3c), but its protein level displayed an opposite pattern to that of MFSD2A, increasing gradually from arteries to veins (Fig. 3d, Supplementary Fig. 3h, Supplementary Fig. 3j). This suggested that CAV1 expression may be regulated by MFSD2A in the DRG vasculature, which is observed at the BBB and and blood-retinal barrier (Andreone et al., 2017; Wang et al., 2020b).

To investigate if caveolar transcytosis was required for macrophage monitoring of DRG endothelium, we used *Cav1*^-/-^ mice and injected BSA and IgG into the tail vein. DRG macrophage uptake of intravenously injected IgG and BSA were both significantly reduced in *Cav1*^-/-^ mice compared to in WT controls (Fig. 3e). When perivascular macrophages were spatially mapped along the arteriovenous axis and analyzed based on which vessel segment they contacted, the largest difference between WT and *Cav1*^-/-^ mice was noted in v-caps (Fig. 3e), which is consistent with the high level of caveolar vesicles observed in this location. This indicated that perivascular macrophages were ingesting material passing across endothelial cells via caveolar vesicles. In support of this hypothesis, using confocal and STED microscopy, we observed that DRG macrophages made direct contact with the cell membrane of CAV1^+^ v-caps (Fig. 3f-g). TEM further confirmed this notion, showing that macrophages and endothelial cells made direct cell-to-cell contact in this location, without being separated by a layer of extracellular matrix or basement membrane (Fig. 3h). Taken together, our data demonstrate a structural zonation across the DRG vasculature that spatially aligns with its permeability. Furthermore, caveolar transcytosis is at least partly required for monitoring of the vasculature by DRG macrophages.

### Two molecularly distinct subsets of macrophages inhabit the DRG

Our data thus far showed that macrophages in the DRG are predominantly perivascular, highly phagocytic and ingest virtually all material crossing through ganglionic blood vessels. These results indicate specialization of the macrophage population at the blood-DRG interface, which prompted us to explore the DRG macrophage pool in greater detail. We used scRNA-seq using the 10x platform to profile the DRG immune landscape at steady state (n=3 mice, 2668 cells). Using dimensionality reduction (UMAP) and unbiased clustering (Louvain) we determined that the DRG was characterized by a heterogenous population of immune cells which included neutrophils (*S100a8, S100a9)*, monocytes (*S100a4, Plac8),* B-cells (*Cd79a*, *Igkc*), T-cells (*Tbrc2, Cd3g),* dendritic cells (DCs) (*Xcr1*), but was numerically dominated by macrophages (*C1qa, Csf1r, Cx3cr1;* 59% of all cells) (Fig. 4a), which separated into two major clusters characterized by *Fcrls, Cd163, Mrc1* (47.2% of macrophages) or *Ccr2 and Cd52* expression (40.2% of macrophages), respectively. Three additional smaller macrophage clusters were present, one displaying interferon-regulated gene expression (*Isg15, Ifi44, Irf7;* 6.4% of macrophages), one expressing stress-induced genes (*Prdx1*, *Ppia*; 5.4% of macrophages) and one cluster expressing a core signature (***T****imd4, **L**yve1, **F**olr2*, TLF) of embryonically-derived self-renewing macrophages (Dick et al., 2022) (0.7% of macrophages).

**Figure 4:**
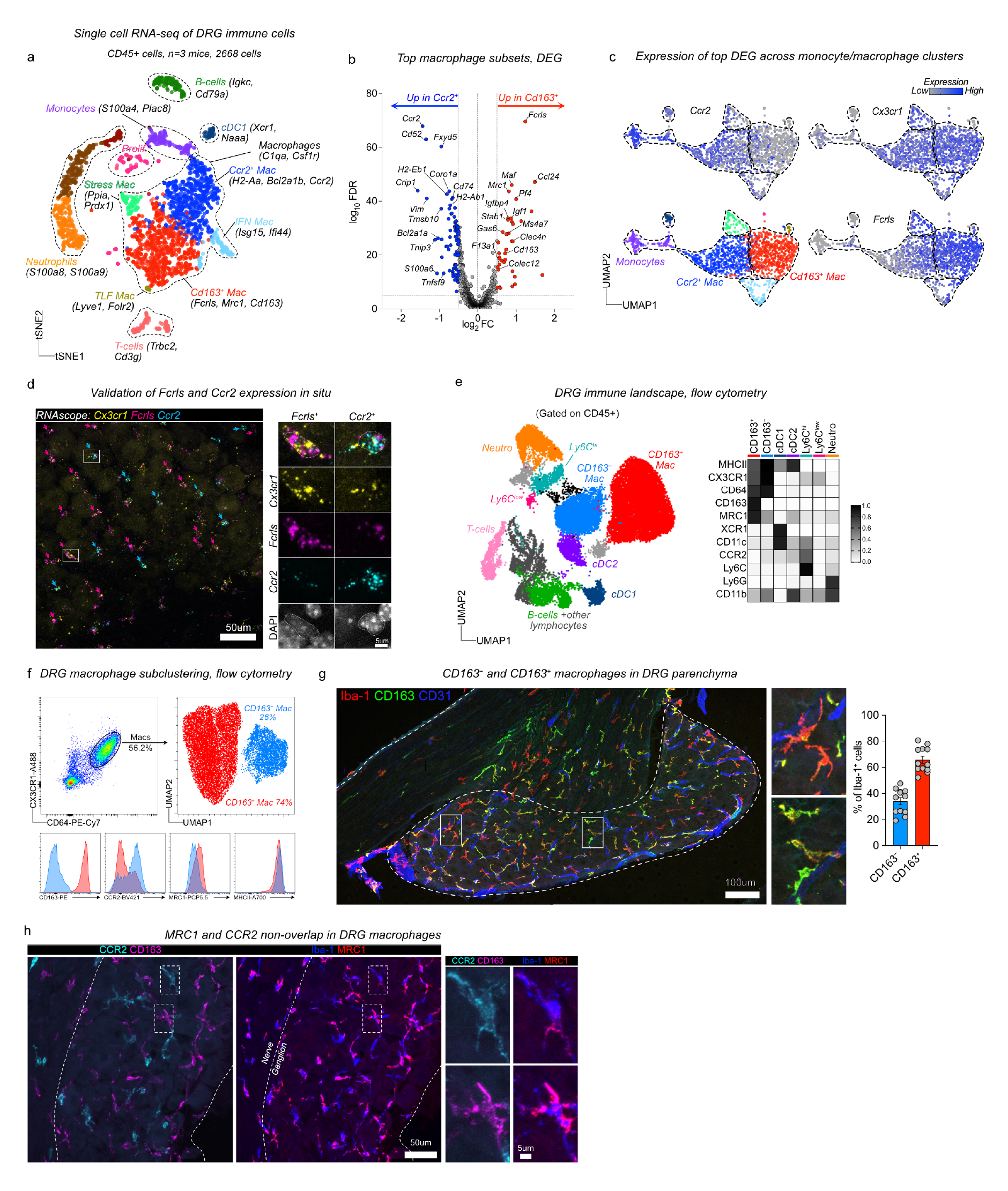
**DRG contains two molecularly distinct macrophage populations** (a) Single-cell RNA-seq analysis of 2668 CD45^+^ DRG cells n= 3 mice. (b) Differential gene expression between CD163^−^ and CD163^+^ macrophages using the Venice algorithm. (c) UMAP of monocyte/macrophage/cDC clusters and their expression of key transcripts. (d) RNAscope of indicated transcripts in DRG sections. Purple arrows indicate *Fcrls*^+^*Ccr2*^−^ cells and turquoise arrows indicate *Fcrls*^−^*Ccr2*^+^ cells. Images are representative of n=3 mice. (e) UMAP of Live, CD45^+^ DRG cells analyzed by flow cytometry. Expression heatmap of selected markers in all myeloid populations. n=4 mice pooled. The experiment was performed three times with similar results. (f) Subclustering of CD64^+^CX3CR1^+^ macrophages from flow cytometry data. (g) Immunostaining and quantification of CD163^−^ and CD163^+^ macrophages in DRG parenchyma. n=12 mice. (h) Immunostaining of MRC1 and CCR2 in CD163^−^ and CD163^+^ macrophages.

We next focused our analysis on the two largest macrophage clusters, as together they made up 87% of the total macrophage pool. Signs of *ex vivo* enzymatic digestion-induced gene expression (Marsh et al., 2022) were apparent, particularly in macrophages (Supplementary Fig. 4a), which included immediate early genes (*Fos, Jun, Atf3, Rhob*). We thus removed these genes after differential gene expression (0.5>log_2_FC; log_10_FDR>5) and before gene ontology analysis. Differential gene expression demonstrated that the largest macrophage cluster (hereafter referred to as CD163^+^ macrophages) highly expressed *Fcrls,* encoding an Fc receptor-like glycoprotein with unknown function, as well as several phagocytic receptors including *Cd163, Mrc1* and *Colec12* (Fig. 4b). Consistently, gene ontology analysis showed enrichment of receptor-mediated endocytosis (Supplementary Table 2). *Maf* was also highly expressed in CD163^+^ macrophages, which is a transcription factor essential for perivascular macrophage survival and function (Silva et al., 2021). *F13a1, Pf4* and *Selenop,* which are all serum factors were also upregulated in CD163^+^ macrophages, again indicating an interaction with the blood. Consistently, platelet degranulation and regulated exocytosis were additional GO-terms associated with CD163^+^ macrophages (Supplementary Table 2). In addition, CD163^+^ macrophages expressed several chemokines (*Ccl7, Ccl8, Ccl12, Ccl24, Pf4*), resulting in enrichment of multiple terms related to chemotaxis (Supplementary Table 2). The second cluster (hereafter referred to as CD163^−^ macrophages) specifically expressed *Ccr2* (Fig 4b-c), a chemokine receptor required for monocyte migration out of the bone marrow (Serbina and Pamer, 2006). Furthermore, *Ccr2* was recently identified as a marker of tissue macrophages that are constantly replaced by circulating monocytes (Dick et al., 2022). 29 ribosomal genes were expressed in this subset, resulting in enrichment of several GO-terms related to protein translation (Supplementary Table 3). After removal of ribosomal genes, GO-terms related to neutrophil functions were enriched (*Lgals3, Adgre5, Anxa2, Tlr2),* as well as mononuclear cell migration and type 2 immune responses (*Lgals3, Tnf, Tnfsf9* and *Cd74)* (Supplementary Table 4). Furthermore, several MHCII genes (*H2-Aa, H2-Ab1, H2- Eb1, H2-DMb1, Cd74*) were upregulated in CD163^−^ macrophages.

We then attempted to validate *in situ* expression of the top two differentially expressed genes (*Fcrls* and *Ccr2,* Fig. 4b-c). Using RNAscope, we confirmed expression of *Fcrls* and *Ccr2* in separate subsets of *Cx3cr1^+^* macrophages, which were both located in the DRG parenchyma, interspersed between neuronal cell bodies (Fig. 4d). In further support of a parenchymal location, we did not detect expression of genes enriched in epineurial sciatic nerve macrophages (*Retnla, Cd209a, Clec10a, Folr2)* (Ydens et al., 2020; Yim et al., 2022) in either of the two DRG macrophage subsets. In fact, several of these genes were highly expressed in the TLF macrophage subset, indicating that this population may in fact correspond to epineurial macrophages (Supplementary Fig. 4b).

We next designed a flow cytometry panel to analyze the DRG myeloid landscape in greater detail. We used dimensionality reduction (UMAP) and unbiased clustering (Phenograph) based on 16 parameters (14 surface antigens, size and granularity) combined with traditional gating (Gating strategy in Supplementary Fig. 4c), and this revealed a similar distribution of macrophages, neutrophils, DCs, monocytes and lymphocytes as for our scRNA-seq data (Fig. 4e, Supplementary Fig. 4d). To investigate macrophage substructure our panel included several pan-macrophage markers (CD11b, CX3CR1, CD64), as well as potential subset-specific antibodies based on our scRNA-seq data (CD163, CCR2, MRC1 and MHCII). We visualized the DRG macrophage pool (CX3CR1^+^CD64^+^) separately, which assigned all macrophages into two major clusters in the resulting UMAP, defined as CD163^+^CCR2^low^ and CD163^−^CCR2^hi^ (Fig. 4f). MRC1 and MHCII expression provided additional separation between these subsets and were more highly expressed by CD163^+^ and CD163^−^macrophages, respectively, which was consistent with our scRNAseq data (Fig. 4f). We next turned to immunohistochemistry and confirmed the presence of CD163^−^Iba1^+^ and CD163^+^Iba1^+^ macrophages with similar frequencies as our flow cytometry data in the DRG tissue parenchyma (Fig. 4g). Consistent with our flow cytometry data we found that when CCR2 and MRC1 antibodies were applied to DRG sections they specifically labeled CD163^−^and CD163^+^ macrophages, respectively (Fig. 4h). In summary, using scRNA-seq, multi-parameter flow cytometry and immunostaining we identified two distinct macrophage subsets in the DRG, best defined by their differential expression of CD163.

### CD163^−^and CD163^+^ macrophages have distinct life cycles

Replenishment of tissue-resident macrophages by circulating monocytes is known to vary across and within tissues (Ginhoux and Guilliams, 2016), shaping macrophage phenotype and function (Blériot et al., 2020). High expression of *Ccr2* was recently identified in a subpopulation of tissue-resident macrophages across several organs that have a high turnover rate from circulating monocytes (Dick et al., 2022). We thus hypothesized that monocytes differentially contribute to the two identified DRG macrophage subsets during steady state, which could have implications on macrophage function. To gain further insight into the relationship between monocytes and CD163^−^ and CD163^+^ macrophages, we removed all lymphocyte, neutrophil and DC clusters from our flow cytometry data and only reclustered cells expressing monocyte or macrophage markers. In the resulting UMAP, Ly6C^hi^ monocytes and CD163^−^ macrophages clustered closely and a new cluster of cells expressing intermediate levels of both monocyte and macrophage markers occupied the space in between (Fig. 5a, referred to as ‘transitioning macrophages’). Ly6C and CCR2 downregulation as well as MHCII, CX3CR1 and CD64 upregulation characterized this transition (Fig. 5b-c), which is consistent with the surface expression changes occurring in ‘monocyte-to-macrophage’ conversion in the intestine (Tamoutounour et al., 2012; Bain et al., 2014). *Ear2* and *Retnla* were recently identified as early genes upregulated in monocytes that have recently infiltrated tissues and are committed to a macrophage fate (Sanin et al., 2022). Consistent with our flow cytometry data, we recorded expression of *Ear2* and *Retnla* in cells situated at the border of the CD163^−^ macrophage and monocyte clusters in the scRNA-seq UMAP (Fig. 5d). In DRG tissue sections, CCR2^+^ cells with monocyte morphology were predominantly situated around the capsule and large veins/venules, suggesting monocyte infiltration and differentiation into CD163^−^CD64^+^CCR2^+^ macrophages occurring at this location (Fig. 5e). Taken together, our flow cytometry and scRNA-seq data indicated that Ly6C^hi^ monocytes replenish CD163^−^ macrophages, while their contribution to CD163^+^ macrophages appears to be limited.

**Figure 5:**
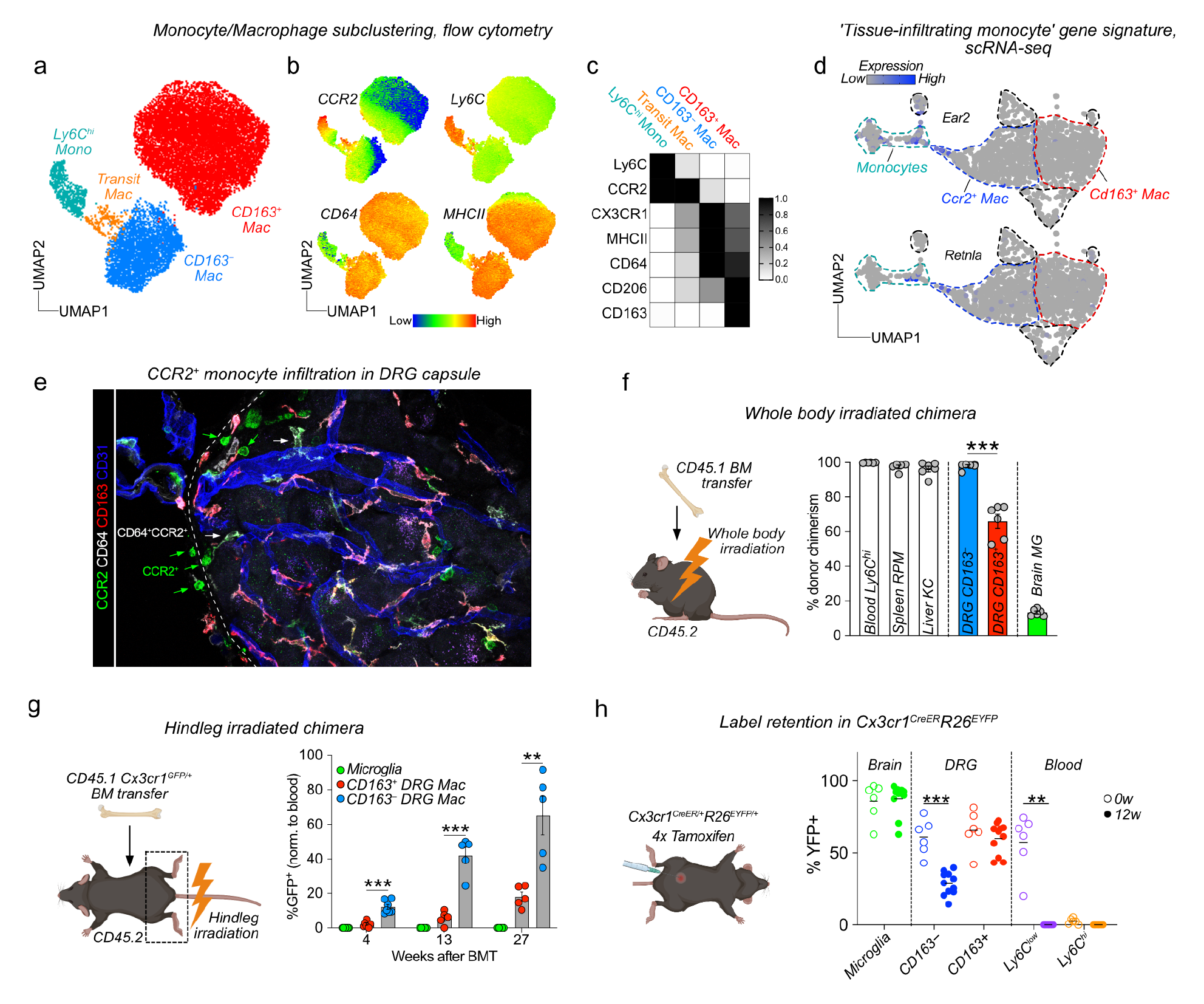
**The two DRG macrophage subsets display different turnover by monocytes** (a) Subclustering of DRG monocyte/macrophage clusters in Fig 4e. n=4 mice pooled. (b) Expression of indicated markers across clusters in (a). (c) Heat map with mean marker expression values for indicated populations. (d) Expression of *Retnla* and *Ear2* (genes expressed in recently infiltrated monocytes, Sanin et al 2022) within monocyte/macrophage clusters. (e) Immunolocalization of CCR2^+^ monocytes and CD64^+^CCR2^+^ macrophages adjacent to the DRG capsule (f) Whole body irradiated BM chimera analyzed 12 weeks after irradiation. Frequency of indicated cell populations that are of donor origin, analyzed by flow cytometry. n=6mice, 1 experiment. Student’s unpaired t-test. (g) CD45.2 mice received hindleg irradiation and given an intravenously injected with 5×10^6^ bone marrow cells from CD45.1:*Cx3cr1*^GFP/+^ mice. Donor chimerism was assessed by analyzing GFP^+^ cells using immunostaining in tissue sections. n=7, 5, 5 mice. 1 experiment/time point. Tukey’s multiple comparisons test. Multiple unpaired t-tests with Holm-Sidak correction. (h) *Cx3cr1*^CreER/+^*R26*^EYFP/+^ mice were given 4×2mg tamoxifen injections i.p. and YFP^+^ cells analyzed in indicated cell populations using flow cytometry after 72h (0w) or 12weeks. (n=6, 11 mice). 2 experiments pooled. Sidak’s multiple comparisons test.

To address experimentally if circulating monocytes differentially contributed to CD163^−^ and CD163^+^ macrophage populations, we turned to bone marrow chimeras. We first lethally irradiated CD45.2 mice and reconstituted them with CD45.1 bone marrow. Analysis of chimerism 12 weeks later revealed complete replacement of circulating Ly6C^hi^ monocytes as well as splenic and liver macrophages (Fig. 5f). Conversely, microglia only displayed 13% replacement by monocyte-derived macrophages, consistent with the well-described radio-resistance of microglia and our own previous data (Lund et al., 2018). In the DRG, CD163^−^ macrophages were completely replaced by monocyte-derived cells and while a majority of CD163^+^ were also donor-derived, 33% remained of host origin (Fig. 5f). This experiment demonstrates that while Ly6C^hi^ monocytes are able to generate both CD163^−^ and CD163^+^ DRG macrophages, CD163^+^ macrophages display partial radio-resistance.

To avoid the macrophage death and tissue inflammation that accompanies whole-body irradiation, we next set up tissue-protected chimeras. We irradiated only the hindlegs of CD45.2 mice and reconstituted them with CD45.1:*Cx3cr1*^GFP/+^ bone marrow (Fig. 5g, Supplementary Fig. 5a), which after 4 weeks resulted in approximately 30% donor chimerism in the blood. We subsequently analyzed macrophage chimerism in the brain and DRGs over several time points up to 26 weeks after irradiation. In contrast to our whole-body chimeras, microglia now remained completely host-derived throughout the study period (Fig. 5g). We next turned to analyzing DRGs, and observed that over time Ly6C^hi^ monocytes differentially contributed to both CD163^−^ and CD163^+^ macrophages: At 4 weeks, CD163^−^ macrophages were 12.1% donor-derived, a number that rose to 42.0% after 13 weeks and 65.1% after 27 weeks (Fig. 5g). For CD163^+^ macrophages these numbers were 2.2% at 4 weeks, 5.7% at 13 weeks and 18.0% at 27 weeks (Fig. 5g). In a parallel set of animals we used flow cytometry on pooled DRGs to confirm the findings at the last time point (Supplementary Fig. 5b).

To validate our results in a setting without irradiation, we made use of *Cx3cr1*^CreER/+^*R26*^EYFP/+^ mice, in which *Cx3cr1* expressing macrophages can be labeled by the administration of tamoxifen. 72 hours after our tamoxifen regimen, on average 85.9% of microglia were YFP^+^, a number that had not changed after 12 weeks (87.6%) (Fig. 5h), which supports the well-described self-sustainability of microglia (Ajami et al., 2007; Ginhoux et al., 2010; Schulz et al., 2012; Hashimoto et al., 2013). Circulating Ly6C^hi^ monocytes displayed negligible labeling at 72 hours and 12 weeks. In the DRG, CD163^−^ and CD163^+^ macrophages displayed equal labeling 72 hours after tamoxifen (61.1 % vs 65.7 % YFP^+^, respectively). 12 weeks later, while CD163^+^ macrophages retained a similar level of YFP expression (60.2%), CD163^−^ macrophages had dropped to 28.5% YFP^+^ (Fig. 5h). Taken together, our data demonstrate that while CD163^−^ macrophages are constantly replenished from circulating monocytes, CD163^+^ macrophages are mostly self-sustained.

### Only CD163^+^ macrophages monitor the vasculature

We next assessed functional differences between the two DRG macrophage subsets. Given that the CD163^+^ macrophages expressed several scavenger receptors and displayed a transcriptional signature associated with endocytosis and interaction with the vasculature, we assessed whether the vasculature monitoring function varied between CD163^−^and CD163^+^ subsets. We first injected fluorescent BSA and analyzed uptake across several organs using flow cytometry. Consistent with our previous analysis of uptake in tissue sections, we found that macrophages in the DRG phagocytosed significantly more BSA than did their ScN or brain counterparts (Fig. 6a). When CD163^−^ and CD163^+^ macrophages were directly compared, uptake was higher in CD163^+^ macrophages (8 fold in the DRG), a finding that was also observed in ScN and the brain (Fig. 6a). We further confirmed a higher phagocytosis of i.v-injected BSA and IgG in CD163^+^ macrophages using DRG tissue sections (Fig. 6b), which additionally allowed us to distinguish macrophages based on their contact with the vasculature. This demonstrated that both CD163 expression (CD163^−^vs CD163^+^) and macrophage contact with the vasculature (perivascular vs parenchymal) determined the level of BSA and IgG uptake (Fig. 6b). Focusing on perivascular CD163^−^and CD163^+^ macrophages, we next performed spatial mapping of macrophage uptake along the arteriovenous axis. Just as we had observed previously, we found that uptake of all tracers peaked in either v-cap or vein-associated macrophages for both subsets (Fig. 6b). Furthermore, this uptake was significantly higher in CD163^+^ compared to CD163^−^macrophages throughout the arteriovenous axis (Fig. 6b). Taken together, our data demonstrate that *in vivo* monitoring of the vasculature is a function restricted to CD163^+^ macrophages.

**Figure 6:**
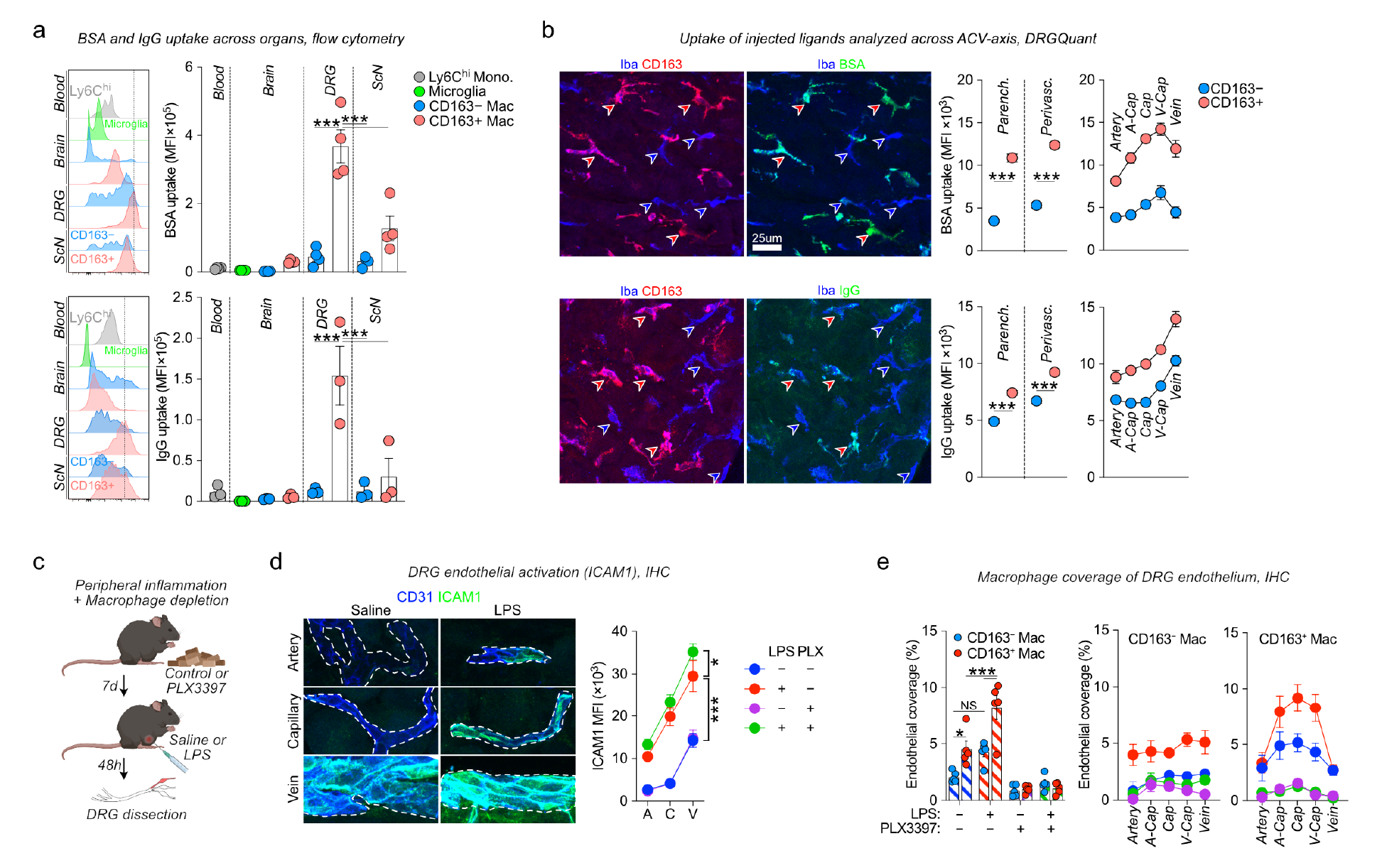
**CD163^+^ macrophages interact functionally with the DRG vasculature** (a) Flow cytometry quantifications of uptake of BSA-A647 (4mg/kg, 1.5h) and IgG-A647 (6mg/kg, 2h) in the indicated organs/cell types. Gated on CD11b^+^Ly6C^hi^ (monocytes), CX3CR1^hi^CD64^low^ (microglia), CX3CR1^+^CD64^hi^ (macrophages). The experiment was performed twice (IgG, n=5 total) or three times (BSA, n=8 total) with similar results. Tukey’s multiple comparisons test. (b) Machine learning (DRGQuant) based analysis of uptake of BSA-A647 (5mg/kg, 1h), goat anti rabbit IgG-A488 (4mg/kg, 4h), 70kDa dextran-TMR (25mg/kg, 1h) in CD163^−^Iba1^+^ and CD163^+^Iba1^+^ macrophages without (parenchymal) or with (perivascular) contact with vasculature. Perivascular macrophages were additionally analyzed based on which vessel segment they contacted. Student’s unpaired t-test. (c) Experiment illustration for macrophage depletion and peripheral endotoxin challenge. Mice were fed chow containing 290 ppm PLX3397 and injected i.p with 1mg/kg LPS. (d) Quantification of ICAM1 expression in CD31^+^ vessel segments. 2-way ANOVA with Tukey’s multiple comparisons test. (e) Quantification of endothelial coverage in DRG sections stained with CD31, Iba-1 and CD163.

### Peripheral inflammation drives arteriovenously zonated activation of endothelium and CD163^+^ macrophages

To further explore macrophage-vasculature interplay in the DRG we next assessed the impact of peripheral inflammation on endothelial cells and macrophages. We injected control or macrophage-depleted mice with LPS or saline and sacrificed them after 48h (Fig. 6c). To measure endothelial activation following LPS injection we performed ICAM1 staining, which demonstrated an arteriovenously zonated expression in all experimental groups. In saline-injected mice, ICAM1 could only be detected in veins. After LPS, ICAM1 staining increased across all vessel segments (Fig. 6d). Macrophages were not required for endothelial activation, as mice fed PLX3397 prior to LPS displayed similar upregulation of ICAM1. We next assessed macrophage contact with the vasculature and recorded that endothelial activation was accompanied by increased coverage of the vasculature by macrophages (Fig. 6e). Mapping of macrophages along the vasculature demonstrated that endothelial coverage was also arteriovenously zonated, peaking in capillaries and that this was largely driven by increased contact with CD163^+^ macrophages (Fig. 6e). These results demonstrate that peripheral inflammation drives zonated activation of both the endothelium and of CD163^+^ macrophages.

### Perivascular CD163^+^ macrophages are conserved in human DRGs

We next addressed whether CD163^−^ and CD163^+^ macrophages were also present in human DRGs. We again made use of a recently published snRNA-seq dataset of human DRGs (Avraham et al., 2022), which contained 2098 macrophages (clusters expressing *CSF1R*). To investigate macrophage substructure we clustered only macrophages, resulting in 5 distinct clusters (Fig 7a). The largest cluster comprised 38.0 % of all macrophages and was characterized by expression of *CD163*, *MRC1, FXIIIA, STAB1* and *COLEC12* (Fig. 7b), thus displaying considerable overlap in gene expression with the CD163^+^ macrophage population identified in mouse DRGs. Gene ontology analysis revealed enrichment of biological processes such as receptor mediated endocytosis (*CD163, COLEC12, TFRC, MRC1, STAB1)*, endothelial tube morphogenesis (*STARD13, RBPJ*) and iron/heme metabolism (*HMOX1, FXIIIA, FTL, BLVRB*) (Supplementary Table 5), indicating that this subset (*CD163*^+^*MRC1*^+^), had a function related to the vasculature, just as we had observed in the mouse. An additional cluster (*CD163*^+^*MRC1*^−^) accounting for 21.2% of macrophages also expressed *CD163* but lacked most of the defining markers of the *CD163*^+^*MRC1*^+^ macrophages, including *FXIIIA*, *COLEC12* and *MRC1.* No genes were specifically upregulated in this cluster. A third cluster accounted for 29.6 % of macrophages and expressed *C3, OXR1* and *KCNIP1* highly, but lacked *CD163*. Differential gene expression also identified *CX3CR1* as a defining marker for this subset (*CD163*^−^*CX3CR1*^+^). While this subset displayed similar frequency as CD163^−^ macrophages in the mouse, it did not overlap in gene expression profile. In addition, two smaller clusters expressing markers related to regulation of neuronal function (CALD1^+^NRXN3^+^, 8.7%) as well as proliferation (TOP2A^+^MKI67^+^, 2.5%) were also evident (Fig. 7a-b). CD163^+^ macrophages are thus conserved in human DRGs and display similar gene expression profiles and putative function.

**Figure 7:**
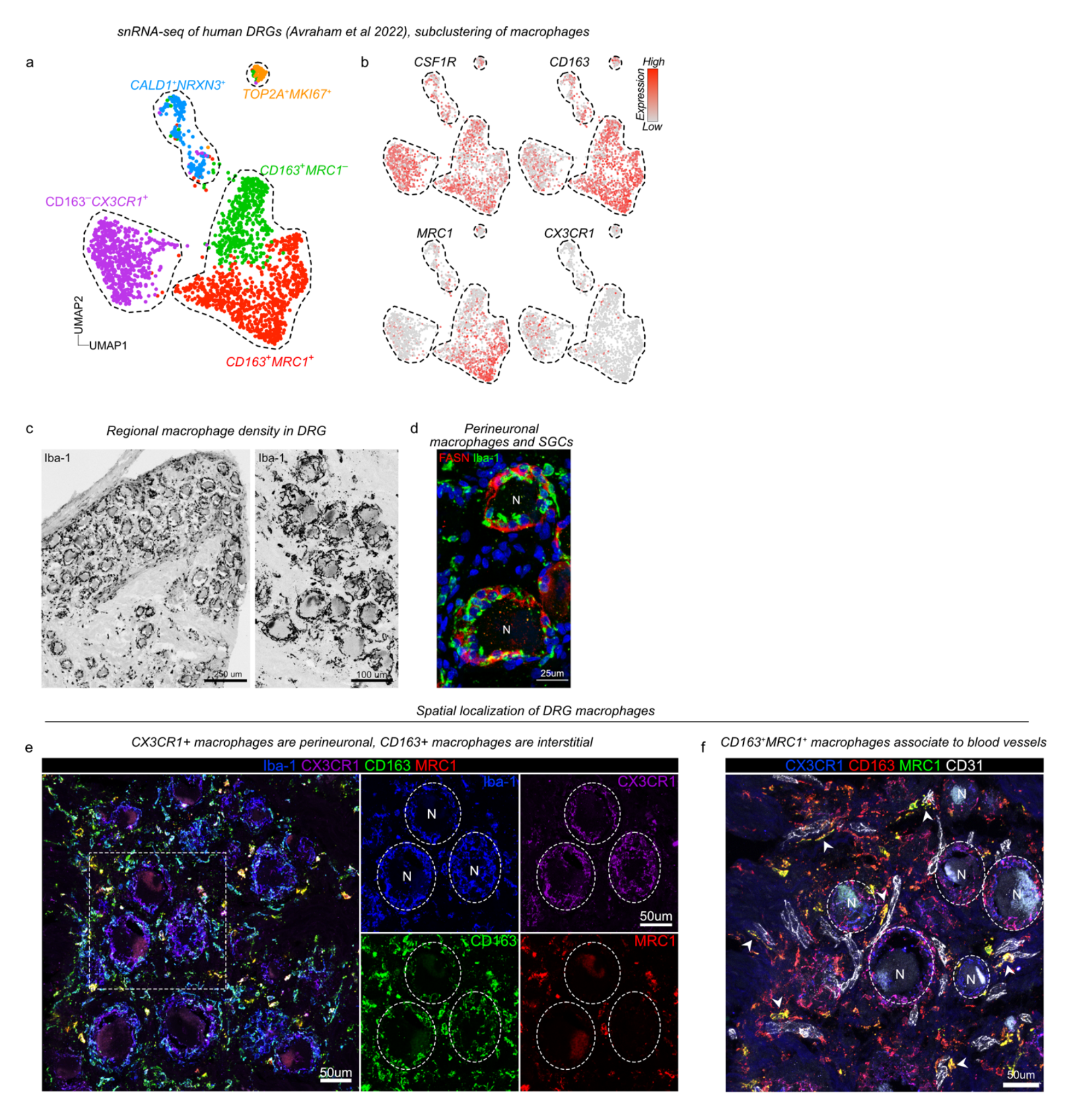
**Perivascular CD163^+^ macrophages are conserved in human DRGs** (a) UMAP of 2098 subclustered macrophage nuclei from 5 human DRG donors (Avraham et al 2022). (b) Expression of key indicated genes across clusters. (c) Iba-1 immunostanining visualizing difference in macrophage distribution between neuron-rich and fiber-rich areas and the presence of perineuronal and ‘interstitial’ Iba-1^+^ macrophages. (d) Immunostaining of Iba-1^+^ perineuronal macrophages and FASN^+^ SGCs, both cell types cluster around neuron cell bodies. (e) Immunostaining showing the perineuronal localization of CX3CR1^+^Iba1^+^ macrophages and the interstitial localization of Iba1^+^ CD163^+^MRC1^+^ macrophages. (f) Immunostaining showing perivascular association of CD163^+^MRC1^+^ macrophages.

In order to validate the presence of the major identified macrophage subsets in human tissues, we again turned to DRG sections from human organ donors (Supplementary Table 1). We first used the pan-macrophage marker Iba-1, which as in the mouse revealed a much higher accumulation of macrophages in the neuronal soma-rich region of DRGs compared to the nerve-rich region (Fig. 7c). In the neuronal soma-rich region macrophages formed dense aggregates around the neuronal cell bodies, but were distinct from FASN^+^ SGCs (Fig. 7d), which also inhabit this space. In addition, macrophages populated the interstitial space, between the neuronal cell bodies. We next applied subset-specific antibodies to DRG sections and observed that CX3CR1 expression was virtually restricted to macrophages surrounding the neuronal soma (Fig. 7e). Conversely, we found that CD163^+^MRC1^+^ and CD163^+^MRC1^−^ macrophages were preferentially located in the interstitium than perineuronally (Fig. 7e). Co- staining of CD31^+^ endothelial cells further revealed that CD163^+^MRC1^+^ were preferentially localized in close contact with endothelial cells (Fig. 7f). In summary, we identify CD163^−^CX3CR1^+^ neuron-associated macrophages which display limited transcriptional and spatial overlap with mouse DRG macrophage subsets. By contrast CD163^+^MRC1^+^ macrophages are transcriptionally and anatomically conserved in human DRGs.

## Discussion

The vasculature is organized into networks of arteries, veins and interconnected capillaries. While this general structure is shared between all organs, transcriptional and anatomical heterogeneity in vascular identity is well characterized (Potente and Mäkinen, 2017; Trimm and Red-Horse, 2022), giving rise to organotypic vascular beds with distinct functional properties. The vasculature of the CNS is characterized by several features to maintain high barrier integrity: low level of transcytosis, constitutive expression of tight and adherens junctions, absence of fenestrae and almost complete coverage of endothelial cells by astrocytes and pericytes (Zhao et al., 2015). The BNB shares several but not all of these features (Malong et al., 2023), and is recognized as the second most restrictive vascular system in the body (Ubogu, 2020). The DRG is highly vascularized, likely as a consequence of the high metabolic demands of DRG sensory neurons. Consistently, DRG neurons and SGCs produce VEGF during development, which is required for proper vascularization of the DRG (Kutcher et al., 2004). In contrast to the BBB and BNB, it is well documented that the blood vessels supplying the sensory ganglia are more permeable to circulating molecules than those supplying the axons (Olsson, 1968; Jacobs et al., 1976; Arvidson, 1979; Kobayashi and Yoshizawa, 2002). However, a cohesive mechanism for this phenomenon has been lacking. We thus undertook a thorough molecular, structural and spatial characterization of the DRG endothelium. Furthermore, the abluminal side of the DRG vasculature was covered with macrophages with what appeared to be an indiscriminate phagocytic ability. We therefore also characterized the hetereogeneity, turnover and function of DRG macrophages.

### Mechanism of blood-ganglion permeability

To address DRG endothelial permeability in an unbiased way we explored the transcriptome of endothelial cells at the single cell level using a recently published dataset (Avraham et al., 2020). This revealed an arteriovenously zonated profile, progressively changing from having BBB-like properties (*Cldn5*^+^) to displaying features of peripheral endothelial beds (*Plvap*^+^). CLDN5 participates in barrier integrity at the BBB and blood-retinal barrier by limiting paracellular diffusion between blood and tissue (Greene et al., 2019). Regional differences in tight junction expression, including CLDN5, between neuronal soma and fiber-rich regions in the DRG have been reported (Hirakawa et al., 2004; Lux et al., 2019). We here clarify that this is a consequence of reduced expression of CLDN5 along the arteriovenous axis, which is a feature shared by several peripheral endothelial beds (Richards et al., 2022), but not the brain vasculature (Vanlandewijck et al., 2018; Kalucka et al., 2020). We did not directly assess the contribution of reduced CLDN5 expression to endothelial permeability, but experimental evidence suggests it is mostly important for small molecular weight compounds. CLDN5 mutant mice display increased permeability only to molecules <0.8kDa at the BBB (Nitta et al., 2003). Outside of the CNS, endothelial cell-specific CLDN5 loss does not affect basal permeability to 10-70 kDa dextrans, but increases histamine-induced permeability in an organ-specific manner (Richards et al., 2022).

Given the high expression of PLVAP in veins we addressed its role in endothelial permeability. PLVAP is restricted to the diaphragm of endothelial fenestrae, transendothelial channels and caveolar vesicles (Stan et al., 1999, 2004). However, the high expression of *Plvap* at mRNA and protein levels in veins and v-caps did not overlap with the absent transendothelial channels and the scarcity of endothelial fenestrae that we observed using TEM and SEM. Several studies have reported endothelial fenestrae in the DRG vasculature (Anzil et al., 1976; Jacobs et al., 1976; Arvidson, 1979; Kobayashi and Yoshizawa, 2002), but their absolute abundance has not been quantified. In venous capillaries, where fenestrae were most abundant, we observed that less than 0.1% of the endothelial lumen was covered with fenestrael openings. By comparison, 12.8% of the luminal surface of the glomerular endothelium of the kidney is covered by fenestrae (Bulger et al., 1983). Based on this finding we find it unlikely that endothelial fenestrae contribute significantly to basal permeability of the DRG endothelium. However, we do not exclude that fenestrae serve other critical purposes important for DRG physiology, such as chemosensation of the body’s internal milieu, similar to sensory circumventricular organs (Miyata, 2015), as previously proposed (Devor, 1999). The limited number of fenestrae and their specific spatial localization to v- caps would support a tightly regulated mechanism.

While fenestrae were limited, as much as 10% of the endothelial cytoplasm in v-caps contained caveolar vesicles. It has been proposed that caveolae and fenestrae are interchangeable structures (Satchell and Braet, 2009). However, we excluded this possibility in the DRG endothelium based on the presence of fenestrae in *Cav1* mutant mice. Furthermore, using these mice we functionally validated the importance of caveolar vesicles for basal permeability of the DRG endothelium. Given that *Cav1* mRNA was not arteriovenously zonated, we speculated that caveolar assembly must be regulated post-translationally. This led us to explore the levels of MFSD2A, a transporter of DHA, an essential fatty acid required for proper brain development and function (Nguyen et al., 2014). The transporter activity of MFSD2A creates a lipid environment in endothelial cells that interferes with caveolar assembly at the BBB (Andreone et al., 2017) and blood-retinal barrier (Wang et al., 2020b). MFSD2A expression is only described in the brain, testis and retinal endothelium. While MFSD2A expression was absent in blood vessels supplying the axons of sensory neurons, it was highly and specifically expressed in ganglionic endothelial cells. In the DRG, MFSD2A expression peaked in a-caps and progressively dropped to non-detectable levels in veins, which negatively correlated with CAV1 expression, thus suggesting MFSD2A could be a negative regulator of caveolar transcytosis also in the DRG. Manipulation of MFSD2A expression at the DRG could thus be a viable strategy to alter blood-ganglion permeability.

Caveolar transcytosis represents a mechanism that could potentially be pharmacologically targeted to reduce DRG endothelial permeability, or alternatively co-opted to deliver drugs into the DRG parenchyma (Kiseleva et al., 2018; Marchetti et al., 2019). This could be desirable in sensory ganglionopathies (Amato and Ropper, 2020), which include paraneoplastic and autoimmune conditions, infections, platinum-based chemotherapy (Dzagnidze et al., 2007; Gupta and Bhaskar, 2016), and likely also fibromyalgia (Goebel et al., 2021; Martínez-Lavín, 2021; Krock et al., 2023).

In addition to the molecular mechanism of DRG endothelial permeability, our findings led us to conclude that the DRG arteriovenous tree is anatomically organized in a conserved manner: arteries and arterial capillaries at the nerve/ganglion interface, venous capillaries directly underlying the capsule, and veins running along the surface of the DRG. Differences in vascularization (Jimenez-Andrade et al., 2008; Kobayashi et al., 2010), expression of tight junctions (Lux et al., 2019) and endothelial permeability between the fiber-rich area and the neuronal soma-rich area of the DRG have been described (Reinhold and Rittner, 2020). Our data now clarifies that this is a consequence of arteries entering the DRG body from the nerve fibers. The impermeable arteries are always localized in the fiber-rich area. By contrast, the neuronal soma-rich area only contains capillaries and veins. Taken together, our data challenges the assumption that the DRG completely lacks an endothelial barrier (Devor, 1999; Haberberger et al., 2019). We instead propose that the DRG endothelial patterning is highly regulated to create a tight barrier in arteries and arterial capillaries which tapers off toward venous capillaries and veins, providing the neuronal soma and interstitial cells underlying the capsule with a high exposure to circulating macromolecules. In addition, we validate CLDN5 and PLVAP as markers of arterial and venous capillaries. This framework can help in the interpretation of future and even previous studies.

### Function of DRG perivascular macrophages

After peripheral nerve damage, macrophages accumulate not only at the site of injury but also around axotomized neuronal cell bodies in the DRG (Niemi et al., 2013; Peng et al., 2016; Kalinski et al., 2020), a process that promotes axon regeneration (Kwon et al., 2013; Niemi et al., 2013; Feng et al., 2023) as well as the development of neuropathic pain (Yu et al., 2020). Most studies of DRG macrophages have been guided by this and analogous findings, which has placed a heavy focus on macrophage-sensory neuron crosstalk in the understanding of DRG macrophage biology (Gheorghe et al., 2022). We here propose that an additional critical function of DRG macrophages is to interact with the vasculature. Several of our experiments support this claim: CD163^+^ macrophages displayed a vasculature-associated transcriptional profile, made close contact with endothelial cells, rapidly phagocytosed circulating macromolecules and increased vessel coverage in response to circulating endotoxin.

CD163^+^ macrophages specifically expressed MRC1 (CD206) at both mRNA and protein levels, which represents a marker that has recently been used to define pain-resolving ‘M2- macrophages’ in DRGs. MRC1^+^ DRG macrophages expand during cisplatin chemotherapy, producing IL-10 that leads to the resolution of pain (Singh et al., 2022). Similarly, in a model of carrageenan-induced hyperalgesia, MRC1^+^ DRG macrophages increase in number and drive pain-resolution by transferring mitochondria to sensory neurons (Vlist et al., 2022). In a model of osteoarthritis pain in which iNOS^+^ macrophages predominate in DRGs and maintain chronic pain, transferring M2-polarized macrophages (expressing MRC1) resolved pain (Raoof et al., 2021). Whether the MRC1^+^ macrophages identified in these reports correspond to the CD163^+^ vasculature-monitoring subset identified in our study remains to be explored. It would also be of particular interest to understand whether these pain-resolving functions are connected with the CD163^+^ macrophage phenotype, and whether it is dependent on contact with or information conveyed through the vasculature.

The observations that DRG macrophages localized around endothelial cells filled with caveolae, and that macrophage phagocytic ability aligned with peak endothelial permeability, leads to the interpretation that the CD163^+^ macrophage network has developed in order to limit enhanced permeability in the DRG. This function is described for perivascular macrophages in the cochlea (Zhang et al., 2012), skin (He et al., 2016) and more recently in the sciatic nerve (Malong et al., 2023). Depletion of macrophages in all these organs results in vessel hyperpermeability. Similarly, CD163^+^ macrophages located in close proximity to fenestrated blood vessels in the area postrema of the brain sequester blood proteins (Willis et al., 2007). However, we did not observe increased leakage of injected tracers into the DRG parenchyma following macrophage depletion. However, we cannot exclude that the non-depleted macrophages (10%) were capable of phagocytosing the injected tracers in the time frame we used in our study (3 hours).

In an effort to link the ontogeny of the DRG macrophage subsets to their phenotype and function (Blériot et al., 2020), we also addressed their turnover from circulating monocytes. This revealed that CD163^+^ macrophages were largely self-sustained, whereas CCR2^+^ macrophages were almost completely replaced by monocytes over 27 weeks. In a recent preprint article, parabiosis experiments confirm the presence of donor-derived macrophages in the DRG parenchyma in naive mice (Guimarães et al., 2022). In addition, our findings agree with the ambitious study by Dick *et al*, which identified tissue-resident CCR2^+^ macrophages in several organs that were almost completely replaced by circulating monocytes (Dick et al., 2022).

Finally, when comparing the transcriptional profile of human and mouse DRG macrophages we concluded that only CD163^+^ macrophages displayed a gene signature that was conserved. Taken together with the perivascular localization of CD163^+^ macrophages in human DRGs, we suggest that CD163^+^ macrophages fulfill an important physiological function associated with endothelial monitoring, as they have been conserved for 90 million years of evolution.

## Methods

### Mice

All mice were either purchased from approved vendors or bred and maintained under specific pathogen-free conditions at Karolinska Institutet, in accordance with national animal care guidelines. All animal experiments were approved by the appropriate ethical review board (Stockholms djurförsöksetiska nämnd). C57Bl/6 mice were purchased from Charles River (C57BL/6J) or bred locally (C57BL/6NTac). BALB/cAnNRj mice were purchased from Janvier. The following strains were originally purchased from the Jackson Laboratory and bred locally: CD45.1 (Jax 002014), *Cav1*^−/−^ mice (Jax 007083) (Razani et al., 2001), *Cx3cr1*^gfp^ (Jax 005582, gift from C. Gerlach, Karolinska Institutet) (Jung et al., 2000), *Cx3cr1*^CreER^ (Jax 020940) (Yona et al., 2013) and R26R-EYFP (Jax 006148) (Srinivas et al., 2001). *Cldn5*(BAC)-eGFP mice (Laviña et al., 2018) were a gift from C. Betsholtz (Uppsala University).

### In vivo studies

#### Macrophage depletion using PLX3397

PLX3397 was formulated into A04 standard diet (Safe Nutrition Service, France) at 75ppm or 290ppm and was administered *ad libitum* for 7 or 21 consecutive days. Identical diet without PLX3397 was used as control.

#### Tamoxifen administration

Tamoxifen (T5648, Sigma) was resuspended in corn oil (C8267, Sigma) and administered i.p to *Cx3cr1*^CreER^*R26*^EYFP^ mice at at 1mg/10g body weight, 4 times over a 5 day period.

#### Whole-body irradiation chimera

CD45.2 mice were irradiated with 9.5 Gray using an X-RAD 320 irradiation source (0.95Gray/minute) with a 20 × 20 cm irradiation field and reconstituted the same day with 5×10^6^ CD45.1 BM cells by tail vein injection. Tissue chimerism was analyzed 12 weeks later.

#### Hindleg bone marrow chimera

Irradiation of only hindlegs was accomplished by maintaining CD45.2 mice under isofluorane anaesthesia and placing the body outside the field of irradiation (the irradiation source is equipped with a lamp to visualize the 20×20 cm irradiation field). Mice were reconstituted the same day with 5×10^6^ BM cells from CD45.1/CD45.2:*Cx3cr1*^gfp/+^ or CD45.1/CD45.1:*Cx3cr1*^gfp/+^ mice by tail vein injection, and analyzed at 4, 13, 26 or 33 weeks.

#### Tracer injections

Anaesthetic cream was applied to the tail 20 min prior to iv injections. Tracers were injected via the tail vein at 4ul/g body weight. Doses and circulation times are summarized in figure legends for each experiment. The following tracers were used: Goat anti Rabbit IgG-A488 (Thermo, A11008), Goat anti Human IgG-A647 (Jackson Immuno, 109-605-003), BSA-A488 (Thermo, A13100), BSA-A647 (Thermo, A34785), 3kDa dextran-TMR (Thermo, D3308), 70kDa dextran-TMR (Thermo, D1818), 500kDa dextran-FITC (Thermo, D7136), 2000kDa dextran-FITC (Thermo, D7137). Mice were subsequently sacrificed and transcardially perfused with PBS only and immediately dissected (flow cytometry analysis) or PBS followed by 4% formaldehyde (immunohistochemistry analysis) followed by 24h post-fixation in 4% formaldehyde.

#### LPS injections

Mice were injected i.p with LPS (1mg/kg) (Condrex, Serotype 0111:B4) or saline vehicle and sacrificed 48h later.

### Whole mount and optical clearing

DRG staining and optical clearing were performed according to (Hunt et al., 2022) using a modified version of the iDISCO protocol (Renier et al., 2014). Briefly, DRGs dissected with the dorsal/ventral roots and peripheral nerve attached were washed and permeabilized in PBS and 0.2% Triton X-100 (PTx) for 3 hours, rinsed in PBS and 0.2% Tween-20 (PTw). DRGs were then incubated 72h with primary antibodies diluted in PTw. After thorough washing in PTw, DRGs were incubated 48h with secondary antibodies diluted in PTw protected from light. Finally, DRGs were washed in PTw before tissue clearing. Stainings were performed in 0.5ml Eppendorf tubes on a tube rotator. DRGs were either imaged directly (CLDN5-GFP) or optically cleared to improve signal depth. Optical clearing was performed in 50ml glass flasks at room temperature (RT) by first dehydrating DRGs in increasing concentrations of tetrahydrofuran (THF) in destilled water (dH_2_O): 50%, 70%; 80%, 2×100% (10 min/solution) followed by refractive index matching in dibenzyl ether (DBE; 2×10min). Cleared tissues were mounted in DBE in custom made 3D-printed image chambers according to (Hunt et al., 2022) and imaged using a confocal microscope.

### Immunostaining

Standard methods for immunostaining were applied. Tissues were collected from animalsperfused with 4% formaldehyde followed by direct dissection and no post-fixation or post-fixation for 24 hours followed by dissection. The different fixation protocols were chosen based on downstream staining protocols. Tissues were embedded in OCT and sectioned using a cryostat at 25-40μm onto SuperFrost Plus glass slides. Sections were stored at −20°C until staining. For stainings, sections were allowed to thaw at RT for 1 hour, washed in PBS for 20 minutes and blocked with 3 % donkey serum in PBS + 0.2% Triton X-100 for 30-60 minutes. Tissues were then incubated with primary antibodies (Supplementary Table 6), diluted in 0.2% Triton X-100, overnight at 4°C. After 3×10 min washes in PBS, secondary antibodies (Thermo or Jackson ImmunoResearch), diluted in PBS, were applied and sections incubated for 2h at RT. After another round of washing, sections were mounted using ProLong gold (Thermo). If not otherwise stated, 3D-images shown in figures were made using Imaris software, 2D-images presented in figures are maximum intensity projections of acquired z-stacks.

### RNAscope

RNAscope was performed according to manufacturer’s protocol for RNAscope Multiplex Fluorescent Detection Reagents v2 (ACD, 323110). Briefly, slides were washed in 1X PBS for 5 min and baked in the HybEZ Oven for 30 min at 60°C. A 15 min fixation in cold 4% PFA was done prior to dehydration with 50%, 70% and twice 100% ethanol for 5 min at RT. Slides were then dried at RT and H_2_O_2_ (ACD, 322335) was applied so that the tissue was covered. This incubation was done for 10 min at RT and the slides were washed in MilliQ water. The slides were transferred into a 1X target retrieval solution (ACD, 322000) at 100°C and kept in this solution for 5 min after which they were washed in MilliQ and 100% ethanol. A circle was drawn around the tissue using a ImmEdge Pen (Vector laboratories, H-4000). The slides were left to dry at RT overnight. Protease III (ACD, 322337) was applied for 30 min at 40°C in the HybEZ Oven. The slides were washed in MilliQ water and a solution of 1X probes was applied for 2 hours at 40°C in the HybEZ Oven. The 1X probe solution was prepared by diluting F*crls*-C2 (ACD, 441231-C2) and *Cx3cr1*-C4 (ACD, 314221-C4) 50 times in 1X *Ccr2*-O1 (ACD, 501681). The slides were washed using 1X wash buffer (ACD, 310091) and the amplification reagents AMP1, AMP2 and AMP3 were applied for 30,30 and 15 minutes respectively at 40°C with washes in wash buffer after each incubation. HRP-C1 was applied for 15 min at 40°C after which the slides were washed in wash buffer and incubated with Opal dye 570 (Akoya, OP-001003) diluted 1:1500 in TSA buffer (ACD, 322809) for 30 min at 40°C. Upon another wash in wash buffer the HRP blocker solution was applied for 15 min at 40°C. The same steps were repeated for HRP-C2 with Opal 690 (Akoya, OP-001006) and HRP-C4 with Opal 520 (Akoya, OP-001001) diluted 1:3000. After the final wash, spectral DAPI was applied to the slides for 30 seconds. The slides were mounted with ProlongGold mounting media.

### Confocal imaging

Z-stack images were acquired using a Zeiss LSM800 laser-scanning confocal microscope equipped with 4 lasers (405nm, 488nm, 561nm and 640nm)

### STED

Super-resolution STED imaging was performed using a STEDYCON (Abberior Instruments) equipped with excitation lasers at 488 nm, 561 nm and 640 nm and a STED laser at 775 nm. Deconvolution was performed on all STED images using Huygens software.

### Transmission electron microscopy (TEM)

Mice were perfused with 2.5 % glutaraldehyde and 1% formaldehyde in 0.1M phosphate buffer, pH 7.4, followed by post-fixation in the same solution (>24h). Following fixation the DRGs were rinsed in 0.1M phosphate buffer prior to post-fixation in 2% osmium tetroxide in 0.1M phosphate buffer, pH 7.4 at 4°C for 2 hour. DRGs were then stepwise dehydrated in ethanol followed by acetone and resin embedded in LX-112 (Ladd). Ultrathin sections (approximately 80-100 nm) were prepared using an EM UC 7 (Leica) and contrasted with uranyl acetate followed by lead citrate. The sections were examined using a Hitachi HT7700 transmission electron microscope (Hitachi High-Technologies) at 80 kV digital images were acquired using a 2kx2k Veleta CCD camera (Olympus Soft Imaging Solutions).

### Scanning Electron Microscopy (SEM)

Following rinsing in 0.1 M phosphate buffer and MilliQ, DRGs were subjected to stepwise dehydration using ethanol and prior to critical point drying in an EM CPD 030 (Leica). The DRG were finally mounted on aluminium pins using double-sided carbon adhesive tabs and platinum coated using a Q150T ES (Quorum). The DRGs were analysed using an Ultra 55 field emission scanning electron microscope (Zeiss) at 3 kV using the SE2 detector.

### Image analysis (Imaris)

Imaris software was used to quantify marker expression across DRG vessel segments. PLVAP and CLDN5-GFP were quantified in sections stained with CD31 and ACTA2 and vessels assigned as arteries (ACTA2^+^, diameter>10um), capillaries (ACTA2^−^, diameter<10um) or veins (ACTA2^+^ on DRG surface, diameter>10um). The zonated expression of PLVAP and CLDN5 in the DRG vasculature was subsequently utilized to quantify MFSD2A and CAV1 expression (MFI) across vessel segments using the following criteria: Arteries (same as above), a-caps (ACTA2^−^, PLVAP^−^ or CLDN5^+^ diameter<10um), v-caps (ACTA2^−^, PLVAP^+^ or CLDN5^−^, diameter<10um). MFI expression was calculated for each vessel segment and normalized to %max for each animal.

### Image analysis (DRGQuant)

Images were analyzed using the DRGquant pipeline described previously (Hunt et al., 2022). In brief, UNET (Ronneberger et al., 2015) models were trained to identify vasculature, macrophages, neuronal-rich regions of the DRG, fiber-rich regions of the DRG, endothelial cells (TEM), vascular lumen (TEM). Model outputs were then run through macros in FIJI (Schindelin et al., 2012) that segmented structures in using connected component analysis in CLIJ (Haase et al., 2020). Macrophages identities were classified as follows: DRG macrophage (≥30% volume within the neuronal soma rich region, Fiber macrophage (Non-DRG macrophage with ≥30% volume in fiber-rich region), parenchymal macrophage (surface area of less than 10μm^2^ in contact with blood vessels), perivascular macrophage (surface area greater than 10μm^2^ in contact with blood vessels). A FIJI macro was written for semi-automated segmentation of the arteriovenous axis with the following workflow: 2D projections of vascular/endothelial cell markers were annotated by an expert as vein, v-cap, capillary, a-cap or artery. Annotations were converted into 2D masks which were then applied to the 3D vasculature map identified via UNET. Perivascular macrophages were then classified based on the vessel type they were most closely associated with. Macrophage morphology as well as fluorescent intensities were quantified using ImageJ’s 3D ROI manager (Ollion et al., 2013). A script was written in python that concatenated all data tables generated in FIJI. For identification of endothelial vesicles (caveolae) in TEM images a stardist (Schmidt et al., 2018) model was trained. A Fiji macro was written that used the UNET output in combination with the stardist output to isolate only vesicles present in the endothelial cells. Single vesicles were defined by a diameter of <100nm and fused vesicles were defined by a diameter greater than 100nm. For all datasets a summary image highlighting all classified objects was generated and visually inspected to ensure the quality of the analysis.

### Generation of single cell suspensions

Mice were euthanized with pentobarbital (APL, 338327) overdose i.p. When applicable, blood was collected from the right ventricle prior to perfusion into an EDTA- or Heparin-coated syringe. Mice were perfused with ice-cold PBS and neural tissues dissected into cold PBS. For analysis by flow cytometry a 5mm coronal slice of the right forebrain was used, both sciatic nerves and approximately 40 pooled cervical, thoracic and lumbar DRGs. Neural tissues were digested in 2mg/ml Collagenase I (Gibco 17100-017), 5mg/ml Dispase II (Sigma, D4693) and 0.5mg/ml DNAse I (Roche, 11284932001) at 37C for 30min (Brain) or 40min (DRGs and ScN). Myelin was removed using 38% Percoll (Sigma, GE17-0891-02) and the cells were subsequently washed and resuspended in PBS. 200 μl blood was lysed in ACK-buffer (Gibco, A1049201) and centrifuged. The pellet was resuspended in PBS and used for staining.

### Flow cytometry

Flow cytometry data was acquired using a Cytek Aurora equipped with violet (405nm), blue (488nm) and red (640nm) lasers. The following antibodies and stains were used: CCR2- BV421 (clone 475301, BD, 747963), Ly6C-BV510 (clone HK1.4, Biolegend, 128033), B220-BV605 (clone RA3-6B2, Biolegend, 103243), Ly6G-BV711 (clone 1A8, Biolegend, 127643), CD11c-BV785 (clone N418, Biolegend, 117336), CX3CR1-A488 (clone SA011F11, Biolegend, 149021), CX3CR1-BV785 (clone SA011F11, Biolegend, 149029), CD163-PE (clone TNKUPJ, Thermo, 12-1631-80), CD64-PE/Cy7 (clone X54-5/7.1, Biolegend, 139313), MRC1-PCP5.5 (clone C068C2, Biolegend, 141716), CD11b-PEFire640 (clone M1/70, Biolegend, 101279) TCRb-APC (clone H57-597, Biolegend, 109212), MHCII-A700 (clone M5/114.15.2, Biolegend, 107622), XCR1-APC/Cy7(clone ZET, Biolegend, 148223), CD45- APC-Fire810 (clone 30F11, Biolegend 103173) CD45.1-A647 (clone A20, Biolegend, 110720), CD45.2- BV785 (clone 104, Biolegend, 109839), LIVE/DEAD Violet dead cell stain kit (Thermo, L34955). Flow cytometry data was analyzed using Flowjo 10 software. Dimensionality reduction and cluster identification was performed using the UMAP and Phenograph packages, respectively.

### Single cell-RNA-seq

Single cell suspensions of DRG cells from BALB/cAnNRj mice were blocked with FcR blocking solution (Miltenyi Biotec l No.130-092-575) and stained with anti-CD45:PE (BioLegend 103105). Additionally, TotalSeq™ anti-mouse Hashtag antibodies were used to label individual samples (BioLegend 155803, 155805, 155809). The samples were strained through a 35-micron filter. DAPI was added to the cell solution in order to exclude dead cells. CD45^+^DAPI^−^ cells were sorted on a BD Influx sorter. The sorted cells were pooled into a single lane on the 10x Genomics Chromium Single Cell 3 ′ v3 system. Library preparation and sequencing were performed at the SciLifeLab sequencing facility, Solna using an Illumina HiSeq 2500 at 61,006 reads/cell. The 10X CellRanger output files (barcodes, feature and count matrix) were analyzed using Seurat in R studio. The filtering criteria was set to include cells with more than 200 and less than 5000 genes as well as less than 5% mitochondrial reads. The data was normalized and scaled using the Seurat functions (NormalizeData and ScalaData). The three samples were demultiplexed using the HTODemux function. Variable features were found using the *vst* method in the FindVariableFeatures function with nfeatures set to 2000 and were used to compute the PCAs using RunPCA. K-nearest and shared nearest neighbor analysis were computed using FindNeighbors with 15 dimensions. Graph-based clustering was conducted using the Louvain algorithm in the FindClusters function with the resolution set at 0.5 (informed by clustertree produced by Clustree). RunUMAP was used to visualize the clusters. Annotation was done using the SingleR package to compare transcriptomes in the ImmunoGen database and based on known cell type markers. For visualization and differential gene expression analysis, data was exported to Bioturing Browser 3.

### Endothelial cell enrichment and qPCR

Single cell suspensions were prepared from liver, lung, kidney, brain and DRGs by enzymatic digestion at 37°C using the following enzymes and incubation times. Liver, small piece of one lobe (1mg/ml Collagenase I Gibco 17100-017, 1mg/ml Collagenase II Wothington LS004176, 5mg/ml Dispase II Sigma D4693 and 7.5ug/ml DNAse I Roche 11284932001, 30min) Lung, one lobe (1mg/ml Collagenase II, 2.5mg/ml Collagenase IV Worthington LS004188 and 15ug/ml DNAse I, 50 min). Kidney, one (2mg/ml Collagenase I and 7.5ug/ml DNAse I, 45min), Forebrain (2mg/ml Collagenase I, 0.5mg/ml DNAse I, 30min) and DRGs, approx. 40 (2mg/ml Collagenase I, 5mg/ml Dispase II, 0.5mg/ml DNAse I, 45min). Single cell suspensions were resuspended in FACS-buffer and labeled with CD31-microbeads (Miltenyi, 130-097-418) and enriched on MS-columns (Miltenyi, 130-042-201) according to manufacturer instructions. Cells were pelleted and lysed in RLT buffer. RNA was isolated using the RNeasy Micro Kit (Qiagen, 74004) and reverse transcribed using iScript cDNA synthesis kit (BioRad, 1708890) according to manufacturer instructions. qPCR was performed on a C1000 Touch thermal cycler equipped with a CFX384 detection module (Bio-Rad) with SYBR Green Master Mix reaction (Bio-Rad, 4367659) and the following PCR primer sequences (Sigma): *Gapdh* (F: TGTAGACCATGTAGTTGAGGTCA, R: AGGTCGGTGTGAACGGATTTG) *Pecam1* F: ACGCTGGTGCTCTATGCAAG, R: TCAGTTGCTGCCCATTCATCA) *Slc2a1* (F: CAGTTCGGCTATAACACTGGTG, R: GCCCCCGACAGAGAAGATG) *Slc7a5* (F: CTTCGGCTCTGTCAATGGGT, R: TTCACCTTGATGGGACGCTC) *Mfsd2a* (F: AAAGACACGCAAAATGCTTACCT, R: AATGAAGGCACAGAGGACGTAGA) *Gpihbp1* (F: AGGGCTGTCCTCCTGATCTTG, R: GGGTCCGCATCACCATCTT) *Slco2a1* (F: ATTAAGGTCTTCGTGCTTTGTCA, R: GTAGGCACTGTAGAGCAACTG).

### Human tissue

Human lumbar dorsal root ganglia were obtained from organ donors through a collaboration with Transplant Quebec. All procedures are approved by and performed in accordance with the ethical review board at McGill University (*McGill University Health Centre REB 2019- 4896).* Familial consent was obtained for each subject. Human DRGs were delivered frozen and prior to use were fixed in 4% formaldehyde for 3-6 hours followed by cryoprotection in 30% sucrose for 3-5 days at 4°C. DRGs were embedded in OCT and sectioned at 12-50μm using a cryostat. Donor details are listed in Supplementary Table 1.

### Analysis of publicly available data

Raw or processed data from the following deposited studies were downloaded and re-clustered and analyzed using commercial code in the Bioturing Browser: Mouse DRG cells, GSE139103 (Avraham et al., 2020). Mouse sciatic nerve macrophages, GSE144707 (Ydens et al., 2020). Mouse multi-organ endothelial cells ArrayExpress: E-MTAB-8077 (Kalucka et al., 2020). Human DRG cells GSE169301 (Avraham et al., 2022). Enrichment scores for BBB and peripheral endothelium specific genes based on bulk RNA-seq of brain vs heart, lung, liver and kidney endothelial cells were retrieved from (Munji et al., 2019). Gene sets for *ex vivo* activation scores were retrieved from (Marsh et al., 2022)

### Funding

This work was supported by The Foundation Blanceflor (to H. Lund), Elisabeth och Alfred Ahlqvists stiftelse (to H. Lund), David och Astrid Hageléns stiftelse (to H. Lund), The Swedish Brain Foundation (to H. Lund), Svenska Sällskapet för Medicinsk Forskning (to H. Lund), Wenner-Gren Foundation (to M. Hunt), Swedish Research Council grant 542-2013- 8373 (to C. I. Svensson) and grant 2013-9306 (to J Lampa), the Knut and Alice Wallenberg Foundation (018.0161 and 2019.0437 to C. I. Svensson), the European Union Seventh Framework, European Union’s Horizon 2020 research and innovation programme under the Marie Skłodowska-Curie Grant Agreement number 764860 (to C. I. Svensson), the European Research Council (ERC) under the European Union’s Horizon 2020 research and innovation programme under the grant agreement no. 866075 (to C. I. Svensson), FOREUM (to C. I. Svensson), a generous donation from Leif Lundblad and family (to C. I. Svensson).

## Acknowledgements

We would like to thank the electron microscopy core facility EMil, Karolinska Institutet, for the TEM and SEM analysis. We acknowledge support for RNA sequencing from the National Genomics Infrastructure in Stockholm funded by Science for Life Laboratory

## References

Ajami B, Bennett JL, Krieger C, Tetzlaff W, Rossi FMV. 2007. Local self-renewal can sustain CNS microglia maintenance and function throughout adult life. Nat Neurosci [Internet] 10:1538–1543. Available from: http://eutils.ncbi.nlm.nih.gov/entrez/eutils/elink.fcgi?dbfrom=pubmed&;id=18026097&retmode=ref&cmd=prlinks

Amato AA, Ropper AH. 2020. Sensory Ganglionopathy. New Engl J Med 383:1657–1662.

Andreone BJ, Chow BW, Tata A, Lacoste B, Ben-Zvi A, Bullock K, Deik AA, Ginty DD, Clish CB, Gu C. 2017. Blood-Brain Barrier Permeability Is Regulated by Lipid Transport-Dependent Suppression of Caveolae-Mediated Transcytosis. Neuron 94:581-594.e5.

Anzil AP, Blinzinger K, Herrlinger H. 1976. Fenestrated blood capillaries in rat cranio-spinal sensory ganglia. Cell Tissue Res 167:563–567.

Armulik A, Genové G, Mäe M, Nisancioglu MH, Wallgard E, Niaudet C, He L, Norlin J, Lindblom P, Strittmatter K, Johansson BR, Betsholtz C. 2010. Pericytes regulate the blood–brain barrier. Nature 468:557–561.

Arvidson B. 1979. Distribution of intravenously injected protein tracers in peripheral ganglia of adult mice. Exp Neurol 63:388–410.

Avraham O, Chamessian A, Feng R, Yang L, Halevi AE, Moore AM, Gereau RW, Cavalli V. 2022. Profiling the molecular signature of satellite glial cells at the single cell level reveals high similarities between rodents and humans. Pain Publish Ahead of Print.

Avraham O, Deng P-Y, Jones S, Kuruvilla R, Semenkovich CF, Klyachko VA, Cavalli V. 2020. Satellite glial cells promote regenerative growth in sensory neurons. Nat Commun 11:4891.

Bain CC, Bravo-Blas A, Scott CL, Perdiguero EG, Geissmann F, Henri S, Malissen B, Osborne LC, Artis D, Mowat AM. 2014. Constant replenishment from circulating monocytes maintains the macrophage pool in the intestine of adult mice. Nat Immunol [Internet] 15:929–937. Available from: http://www.nature.com/doifinder/10.1038/ni.2967

Blériot C, Chakarov S, Ginhoux F. 2020. Determinants of Resident Tissue Macrophage Identity and Function. Immunity 52:957–970.

Bonnardel J, T’Jonck W, Gaublomme D, Browaeys R, Scott CL, Martens L, Vanneste B, Prijck SD, Nedospasov SA, Kremer A, Hamme EV, Borghgraef P, Toussaint W, Bleser PD, Mannaerts I, Beschin A, Grunsven LA van, Lambrecht BN, Taghon T, Lippens S, Elewaut D, Saeys Y, Guilliams M. 2019. Stellate Cells, Hepatocytes, and Endothelial Cells Imprint the Kupffer Cell Identity on Monocytes Colonizing the Liver Macrophage Niche. Immunity [Internet] 51:638–654.e9. Available from: https://linkinghub.elsevier.com/retrieve/pii/S1074761319303681

Bulger RE, Eknoyan G, Purcell DJ, Dobyan DC. 1983. Endothelial characteristics of glomerular capillaries in normal, mercuric chloride-induced, and gentamicin-induced acute renal failure in the rat. J Clin Invest 72:128–141.

Chen L, Hua K, Zhang N, Wang J, Meng J, Hu Z, Gao H, Li F, Chen Y, Ren J, Liu L, Zhou Q, Gu J, Song J, Zhang X, Hu S. 2022. Multifaceted Spatial and Functional Zonation of Cardiac Cells in Adult Human Heart. Circulation 145:315–318.

Devor M. 1999. Unexplained peculiarities of the dorsal root ganglion. Pain 82:S27–S35.

Dick SA, Wong A, Hamidzada H, Nejat S, Nechanitzky R, Vohra S, Mueller B, Zaman R, Kantores C, Aronoff L, Momen A, Nechanitzky D, Li WY, Ramachandran P, Crome SQ, Becher B, Cybulsky MI, Billia F, Keshavjee S, Mital S, Robbins CS, Mak TW, Epelman S. 2022. Three tissue resident macrophage subsets coexist across organs with conserved origins and life cycles. Sci Immunol 7:eabf7777.

Dzagnidze A, Katsarava Z, Makhalova J, Liedert B, Yoon M-S, Kaube H, Limmroth V, Thomale J. 2007. Repair Capacity for Platinum-DNA Adducts Determines the Severity of Cisplatin-Induced Peripheral Neuropathy. J Neurosci 27:9451–9457.

Feng R, Saraswathy VM, Mokalled MH, Cavalli V. 2023. Self-renewing macrophages in dorsal root ganglia contribute to promote nerve regeneration. Proc National Acad Sci 120:e2215906120.

Gheorghe R-O, Grosu AV, Bica-Popi M, Ristoiu V. 2022. The Yin/Yang Balance of Communication between Sensory Neurons and Macrophages in Traumatic Peripheral Neuropathic Pain. Int J Mol Sci 23:12389.

Ginhoux F, Greter M, Leboeuf M, Nandi S, See P, Gokhan S, Mehler MF, Conway SJ, Ng LG, Stanley ER, Samokhvalov IM, Merad M. 2010. Fate Mapping Analysis Reveals That Adult Microglia Derive from Primitive Macrophages. Science [Internet] 330:841–845. Available from: http://www.sciencemag.org/cgi/doi/10.1126/science.1194637

Ginhoux F, Guilliams M. 2016. Tissue-Resident Macrophage Ontogeny and Homeostasis. Immunity [Internet] 44:439–449. Available from: http://linkinghub.elsevier.com/retrieve/pii/S1074761316300632

Godel T, Pham M, Heiland S, Bendszus M, Bäumer P. 2016. Human dorsal-root-ganglion perfusion measured in-vivo by MRI. Neuroimage [Internet] 141:81–87. Available from: https://linkinghub.elsevier.com/retrieve/pii/S1053811916303366

Goebel A, Krock E, Gentry C, Israel MR, Jurczak A, Urbina CM, Sandor K, Vastani N, Maurer M, Cuhadar U, Sensi S, Nomura Y, Menezes J, Baharpoor A, Brieskorn L, Sandström A, Tour J, Kadetoff D, Haglund L, Kosek E, Bevan S, Svensson CI, Andersson DA. 2021. Passive transfer of fibromyalgia symptoms from patients to mice. J Clin Invest 131:e144201.

Gosselin D, Link VM, Romanoski CE, Fonseca GJ, Eichenfield DZ, Spann NJ, Stender JD, Chun HB, Garner H, Geissmann F, Glass CK. 2014. Environment drives selection and function of enhancers controlling tissue-specific macrophage identities. Cell [Internet] 159:1327–1340. Available from: http://linkinghub.elsevier.com/retrieve/pii/S0092867414015001

Greene C, Hanley N, Campbell M. 2019. Claudin-5: gatekeeper of neurological function. Fluids Barriers Cns 16:3.

Guilliams M, Thierry GR, Bonnardel J, Bajenoff M. 2020. Establishment and Maintenance of the Macrophage Niche. Immunity 52:434–451.

Guimarães RM, Silva CEA da, Davoli-Ferreira M, Gomes FIF, Mendes A, Fonseca MM, Damasceno S, Andrade LP, Cunha FQ, Alves-Filho JC, Cunha TM. 2022. Sensory neuron-associated macrophages proliferate in the sensory ganglia after peripheral nerve injury in a CX3CR1 signaling dependent manner. Biorxiv:2022.03.22.485276.

Guo L, Zhang H, Hou Y, Wei T, Liu J. 2016. Plasmalemma vesicle-associated protein: A crucial component of vascular homeostasis. Exp Ther Med 12:1639–1644.

Gupta R, Bhaskar A. 2016. Chemotherapy-induced peripheral neuropathic pain. Bja Educ 16:115–119.

Haase R, Royer LA, Steinbach P, Schmidt D, Dibrov A, Schmidt U, Weigert M, Maghelli N, Tomancak P, Jug F, Myers EW. 2020. CLIJ: GPU-accelerated image processing for everyone. Nat Methods 17:5–6.

Haberberger RV, Barry C, Dominguez N, Matusica D. 2019. Human Dorsal Root Ganglia. Front Cell Neurosci 13:271.

Hanani M, Spray DC. 2020. Emerging importance of satellite glia in nervous system function and dysfunction. Nat Rev Neurosci 21:485–498.

Hashimoto D, Chow A, Noizat C, Teo P, Beasley MB, Leboeuf M, Becker CD, See P, Price J, Lucas D, Greter M, Mortha A, Boyer SW, Forsberg EC, Tanaka M, Rooijen NV, García- Sastre A, Stanley ER, Ginhoux F, Frenette PS, Merad M. 2013. Tissue-Resident Macrophages Self-Maintain Locally throughout Adult Life with Minimal Contribution from Circulating Monocytes. Immunity [Internet] 38:792–804. Available from: http://linkinghub.elsevier.com/retrieve/pii/S107476131300157X

He H, Mack JJ, c EG, Warren CM, Squadrito ML, Kilarski WW, Baer C, Freshman RD, McDonald AI, Ziyad S, Swartz MA, Palma MD, Iruela-Arispe ML. 2016. Perivascular Macrophages Limit Permeability. Arteriosclerosis Thrombosis Vasc Biology [Internet] 36:2203–2212. Available from: https://www.ahajournals.org/doi/10.1161/ATVBAHA.116.307592

Hirakawa H, Okajima S, Nagaoka T, Kubo T, Takamatsu T, Oyamada M. 2004. Regional differences in blood–nerve barrier function and tight-junction protein expression within the rat dorsal root ganglion. Neuroreport 15:405–408.

Hoeffel G, Chen J, Lavin Y, Low D, Almeida FF, See P, Beaudin AE, Lum J, Low I, Forsberg EC, Poidinger M, Zolezzi F, Larbi A, Ng LG, Chan JKY, Greter M, Becher B, Samokhvalov IM, Merad M, Ginhoux F. 2015. C-Myb+ Erythro-Myeloid Progenitor-Derived Fetal Monocytes Give Rise to Adult Tissue-Resident Macrophages. Immunity [Internet] 42:665–678. Available from: http://linkinghub.elsevier.com/retrieve/pii/S1074761315001259

Hunt MA, Lund H, Delay L, Santos GGD, Pham A, Kurtovic Z, Telang A, Lee A, Parvathaneni A, Kussick E, Corr M, Yaksh TL. 2022. DRGquant: A new modular AI- based pipeline for 3D analysis of the DRG. J Neurosci Meth 371:109497.

Jacobs JM, Macfarlane RM, Cavanagh JB. 1976. Vascular leakage in the dorsal root ganglia of the rat, studied with horseradish peroxidase. J Neurol Sci [Internet] 29:95–107. Available from: https://linkinghub.elsevier.com/retrieve/pii/0022510X76900836

Jimenez-Andrade JM, Herrera MB, Ghilardi JR, Vardanyan M, Melemedjian OK, Mantyh PW. 2008. Vascularization of the dorsal root ganglia and peripheral nerve of the mouse: implications for chemical-induced peripheral sensory neuropathies. Mol Pain [Internet] 4:10. Available from: http://journals.sagepub.com/doi/10.1186/1744-8069-4-10

Jung S, Aliberti J, Graemmel P, Sunshine MJ, Kreutzberg GW, Sher A, Littman DR. 2000. Analysis of Fractalkine Receptor CX 3 CR1 Function by Targeted Deletion and Green Fluorescent Protein Reporter Gene Insertion. Mol Cell Biol 20:4106–4114.

Kalinski AL, Yoon C, Huffman LD, Duncker PC, Kohen R, Passino R, Hafner H, Johnson C, Kawaguchi R, Carbajal KS, Jara JS, Hollis E, Geschwind DH, Segal BM, Giger RJ. 2020. Analysis of the immune response to sciatic nerve injury identifies efferocytosis as a key mechanism of nerve debridement. Elife 9:e60223.

Kalucka J, Rooij LPMH de, Goveia J, Rohlenova K, Dumas SJ, Meta E, Conchinha NV, Taverna F, Teuwen L-A, Veys K, García-Caballero M, Khan S, Geldhof V, Sokol L, Chen R, Treps L, Borri M, Zeeuw P de, Dubois C, Karakach TK, Falkenberg KD, Parys M, Yin X, Vinckier S, Du Y, Fenton RA, Schoonjans L, Dewerchin M, Eelen G, Thienpont B, Lin L, Bolund L, Li X, Luo Y, Carmeliet P. 2020. Single-Cell Transcriptome Atlas of Murine Endothelial Cells. Cell [Internet] 180:764–779.e20. Available from: https://linkinghub.elsevier.com/retrieve/pii/S0092867420300623

Kiseleva RYu, Glassman PM, Greineder CF, Hood ED, Shuvaev VV, Muzykantov VR. 2018. Targeting therapeutics to endothelium: are we there yet? Drug Deliv Transl Re 8:883–902.

Kobayashi S, Mwaka ES, Baba H, Takeno K, Miyazaki T, Matsuo H, Uchida K, Meir A. 2010. Microvascular system of the lumbar dorsal root ganglia in rats. Part I: a 3D analysis with scanning electron microscopy of vascular corrosion casts: Laboratory investigation. J Neurosurg Spine 12:197–202.

Kobayashi S, Yoshizawa H. 2002. Effect of mechanical compression on the vascular permeability of the dorsal root ganglion. J Orthopaed Res 20:730–739.

Kolter J, Kierdorf K, Henneke P. 2020. Origin and Differentiation of Nerve-Associated Macrophages. J Immunol 204:271–279.

Krock E, Morado-Urbina CE, Menezes J, Hunt MA, Sandström A, Kadetoff D, Tour J, Verma V, Kultima K, Haglund L, Meloto CB, Diatchenko L, Kosek E, Svensson CI. 2023. Fibromyalgia patients with elevated levels of anti–satellite glia cell immunoglobulin G antibodies present with more severe symptoms. Pain Publish Ahead of Print.

Kutcher ME, Klagsbrun M, Mamluk R. 2004. VEGF is required for the maintenance of dorsal root ganglia blood vessels but not neurons during development. Faseb J 18:1952–1954.

Kwon MJ, Kim J, Shin H, Jeong SR, Kang YM, Choi JY, Hwang DH, Kim BG. 2013. Contribution of macrophages to enhanced regenerative capacity of dorsal root ganglia sensory neurons by conditioning injury. J Neurosci [Internet] 33:15095–15108. Available from: http://www.jneurosci.org/cgi/doi/10.1523/JNEUROSCI.0278-13.2013

Lavin Y, Winter D, Blecher-Gonen R, David E, Keren-Shaul H, Merad M, Jung S, Amit I. 2014. Tissue-resident macrophage enhancer landscapes are shaped by the local microenvironment. Cell [Internet] 159:1312–1326. Available from: http://linkinghub.elsevier.com/retrieve/pii/S0092867414014494

Laviña B, Castro M, Niaudet C, Cruys B, Álvarez-Aznar A, Carmeliet P, Bentley K, Brakebusch C, Betsholtz C, Gaengel K. 2018. Defective endothelial cell migration in the absence of Cdc42 leads to capillary-venous malformations. Development 145:dev161182.

Lund H, Pieber M, Parsa R, Han J, Grommisch D, Ewing E, Kular L, Needhamsen M, Espinosa A, Nilsson E, Överby AK, Butovsky O, Jagodic M, Zhang X-M, Harris RA. 2018. Competitive repopulation of an empty microglial niche yields functionally distinct subsets of microglia-like cells. Nat Commun [Internet] 9:4845. Available from: http://www.nature.com/articles/s41467-018-07295-7

Lux TJ, Hu X, Ben-Kraiem A, Blum R, Chen JT-C, Rittner HL. 2019. Regional Differences in Tight Junction Protein Expression in the Blood–DRG Barrier and Their Alterations after Nerve Traumatic Injury in Rats. Int J Mol Sci 21:270.

Malong L, Napoli I, Casal G, White IJ, Stierli S, Vaughan A, Cattin A-L, Burden JJ, Hng KI, Bossio A, Flanagan A, Zhao HT, Lloyd AC. 2023. Characterization of the structure and control of the blood-nerve barrier identifies avenues for therapeutic delivery. Dev Cell 58:174–191.e8.

Marchetti GM, Burwell TJ, Peterson NC, Cann JA, Hanna RN, Li Q, Ongstad EL, Boyd JT, Kennedy MA, Zhao W, Rickert KW, Grimsby JS, Dall’Acqua WF, Wu H, Tsui P, Borrok MJ, Gupta R. 2019. Targeted drug delivery via caveolae-associated protein PV1 improves lung fibrosis. Commun Biology 2:92.

Marsh SE, Walker AJ, Kamath T, Dissing-Olesen L, Hammond TR, Soysa TY de, Young AMH, Murphy S, Abdulraouf A, Nadaf N, Dufort C, Walker AC, Lucca LE, Kozareva V, Vanderburg C, Hong S, Bulstrode H, Hutchinson PJ, Gaffney DJ, Hafler DA, Franklin RJM, Macosko EZ, Stevens B. 2022. Dissection of artifactual and confounding glial signatures by single-cell sequencing of mouse and human brain. Nat Neurosci 25:306–316.

Martínez-Lavín M. 2021. Dorsal root ganglia: fibromyalgia pain factory? Clin Rheumatol 40:783–787.

Mass E, Nimmerjahn F, Kierdorf K, Schlitzer A. 2023. Tissue-specific macrophages: how they develop and choreograph tissue biology. Nat Rev Immunol:1–17.

Mildenberger W, Stifter SA, Greter M. 2022. Diversity and function of brain-associated macrophages. Curr Opin Immunol 76:102181.

Miyata S. 2015. New aspects in fenestrated capillary and tissue dynamics in the sensory circumventricular organs of adult brains. Front Neurosci-switz 9:390.

Molawi K, Wolf Y, Kandalla PK, Favret J, Hagemeyer N, Frenzel K, Pinto AR, Klapproth K, Henri S, Malissen B, Rodewald H-R, Rosenthal NA, Bajenoff M, Prinz M, Jung S, Sieweke MH. 2014. Progressive replacement of embryo-derived cardiac macrophages with age. J Exp Medicine [Internet] 211:2151–2158. Available from: http://eutils.ncbi.nlm.nih.gov/entrez/eutils/elink.fcgi?dbfrom=pubmed&;id=25245760&retmode=ref&cmd=prlinks

Munji RN, Soung AL, Weiner GA, Sohet F, Semple BD, Trivedi A, Gimlin K, Kotoda M, Korai M, Aydin S, Batugal A, Cabangcala AC, Schupp PG, Oldham MC, Hashimoto T, Noble-Haeusslein LJ, Daneman R. 2019. Profiling the mouse brain endothelial transcriptome in health and disease models reveals a core blood-brain barrier dysfunction module. Nat Neurosci [Internet] 22:1892–1902. Available from: http://www.nature.com/articles/s41593-019-0497-x

Nguyen LN, Ma D, Shui G, Wong P, Cazenave-Gassiot A, Zhang X, Wenk MR, Goh ELK, Silver DL. 2014. Mfsd2a is a transporter for the essential omega-3 fatty acid docosahexaenoic acid. Nature 509:503–506.

Niemi JP, DeFrancesco-Lisowitz A, Roldán-Herńandez L, Lindborg JA, Mandell D, Zigmond RE. 2013. A critical role for macrophages near axotomized neuronal cell bodies in stimulating nerve regeneration. J Neurosci [Internet] 33:16236–16248. Available from: http://www.jneurosci.org/cgi/doi/10.1523/JNEUROSCI.3319-12.2013

Nitta T, Hata M, Gotoh S, Seo Y, Sasaki H, Hashimoto N, Furuse M, Tsukita S. 2003. Size-selective loosening of the blood-brain barrier in claudin-5–deficient mice. J Cell Biology 161:653–660.

Ollion J, Cochennec J, Loll F, Escudé C, Boudier T. 2013. TANGO: a generic tool for high-throughput 3D image analysis for studying nuclear organization. Bioinformatics 29:1840– 1841.

Olsson Y. 1968. Topographical differences in the vascular permeability of the peripheral nervous system. Acta Neuropathol [Internet] 10:26–33. Available from: http://link.springer.com/10.1007/BF00690507

Park MD, Silvin A, Ginhoux F, Merad M. 2022. Macrophages in health and disease. Cell 185:4259–4279.

Parton RG, McMahon K-A, Wu Y. 2020. Caveolae: Formation, dynamics, and function. Curr Opin Cell Biol 65:8–16.

Peng J, Gu N, Zhou L, Eyo UB, Murugan M, Gan W-B, Wu L-J. 2016. Microglia and monocytes synergistically promote the transition from acute to chronic pain after nerve injury. Nat Commun [Internet] 7:12029. Available from: http://www.nature.com/doifinder/10.1038/ncomms12029

Perdiguero EG, Klapproth K, Schulz C, Busch K, Azzoni E, Crozet L, Garner H, Trouillet C, Bruijn MF de, Geissmann F, Rodewald H-R. 2015. Tissue-resident macrophages originate from yolk-sac-derived erythro-myeloid progenitors. Nature [Internet] 518:547–551. Available from: http://www.nature.com/doifinder/10.1038/nature13989

Potente M, Mäkinen T. 2017. Vascular heterogeneity and specialization in development and disease. Nat Rev Mol Cell Bio 18:477–494.

Profaci CP, Munji RN, Pulido RS, Daneman R. 2020. The blood–brain barrier in health and disease: Important unanswered questions. J Exp Med 217:e20190062.

Raoof R, Gil CM, Lafeber FPJG, Visser H de, Prado J, Versteeg S, Pascha MN, Heinemans ALP, Adolfs Y, Pasterkamp J, Wood JN, Mastbergen SC, Eijkelkamp N. 2021. Dorsal Root Ganglia Macrophages Maintain Osteoarthritis Pain. J Neurosci 41:8249–8261.

Razani B, Engelman JA, Wang XB, Schubert W, Zhang XL, Marks CB, Macaluso F, Russell RG, Li M, Pestell RG, Vizio DD, Hou H, Kneitz B, Lagaud G, Christ GJ, Edelmann W, Lisanti MP. 2001. Caveolin-1 Null Mice Are Viable but Show Evidence of Hyperproliferative and Vascular Abnormalities*. J Biol Chem 276:38121–38138.

Reinhold AK, Rittner HL. 2020. Characteristics of the nerve barrier and the blood dorsal root ganglion barrier in health and disease. Exp Neurol 327:113244.

Renier N, Wu Z, Simon DJ, Yang J, Ariel P, Tessier-Lavigne M. 2014. iDISCO: A Simple, Rapid Method to Immunolabel Large Tissue Samples for Volume Imaging. Cell 159:896– 910.

Richards M, Nwadozi E, Pal S, Martinsson P, Kaakinen M, Gloger M, Sjöberg E, Koltowska K, Betsholtz C, Eklund L, Nordling S, Claesson-Welsh L. 2022. Claudin5 protects the peripheral endothelial barrier in an organ and vessel-type-specific manner. Elife 11:e78517.

Richner M, Ferreira N, Dudele A, Jensen TS, Vaegter CB, Gonçalves NP. 2019. Functional and Structural Changes of the Blood-Nerve-Barrier in Diabetic Neuropathy. Front Neurosci-switz 12:1038.

Ronneberger O, Fischer P, Brox T. 2015. U-Net: Convolutional Networks for Biomedical Image Segmentation. Arxiv.

Sanin DE, Ge Y, Marinkovic E, Kabat AM, Castoldi A, Caputa G, Grzes KM, Curtis JD, Thompson EA, Willenborg S, Dichtl S, Reinhardt S, Dahl A, Pearce EL, Eming SA, Gerbaulet A, Roers A, Murray PJ, Pearce EJ. 2022. A common framework of monocyte-derived macrophage activation. Sci Immunol 7:eabl7482.

Satchell SC, Braet F. 2009. Glomerular endothelial cell fenestrations: an integral component of the glomerular filtration barrier. Am J Physiol-renal 296:F947–F956.

Schindelin J, Arganda-Carreras I, Frise E, Kaynig V, Longair M, Pietzsch T, Preibisch S, Rueden C, Saalfeld S, Schmid B, Tinevez J-Y, White DJ, Hartenstein V, Eliceiri K, Tomancak P, Cardona A. 2012. Fiji: an open-source platform for biological-image analysis. Nat Methods 9:676–682.

Schmidt U, Weigert M, Broaddus C, Myers G. 2018. Medical Image Computing and Computer Assisted Intervention – MICCAI 2018, 21st International Conference, Granada, Spain, September 16-20, 2018, Proceedings, Part II. Lect Notes Comput Sc:265–273.

Schulz C, Perdiguero EG, Chorro L, Szabo-Rogers H, Cagnard N, Kierdorf K, Prinz M, Wu B, Jacobsen SEW, Pollard JW, Frampton J, Liu KJ, Geissmann F. 2012. A Lineage of Myeloid Cells Independent of Myb and Hematopoietic Stem Cells. Science [Internet] 336:86–90. Available from: http://www.sciencemag.org/cgi/doi/10.1126/science.1219179

Serbina NV, Pamer EG. 2006. Monocyte emigration from bone marrow during bacterial infection requires signals mediated by chemokine receptor CCR2. Nat Immunol [Internet] 7:311–317. Available from: http://www.nature.com/doifinder/10.1038/ni1309

Shaw TN, Houston SA, Wemyss K, Bridgeman HM, Barbera TA, Zangerle-Murray T, Strangward P, Ridley AJL, Wang P, Tamoutounour S, Allen JE, Konkel JE, Grainger JR. 2018. Tissue-resident macrophages in the intestine are long lived and defined by Tim-4 and CD4 expression. J Exp Med 215:1507–1518.

Silva HM, Kitoko JZ, Queiroz CP, Kroehling L, Matheis F, Yang KL, Reis BS, Ren-Fielding C, Littman DR, Bozza MT, Mucida D, Lafaille JJ. 2021. c-MAF–dependent perivascular macrophages regulate diet-induced metabolic syndrome. Sci Immunol 6:eabg7506.

Singh SK, Krukowski K, Laumet GO, Weis D, Alexander JF, Heijnen CJ, Kavelaars A. 2022. CD8+ T cell–derived IL-13 increases macrophage IL-10 to resolve neuropathic pain. Jci Insight 7:e154194.

Srinivas S, Watanabe T, Lin C-S, William CM, Tanabe Y, Jessell TM, Costantini F. 2001. Cre reporter strains produced by targeted insertion of EYFP and ECFP into the ROSA26 locus. Bmc Dev Biol 1:4.

Stan R-V, Kubitza M, Palade GE. 1999. PV-1 is a component of the fenestral and stomatal diaphragms in fenestrated endothelia. Proc National Acad Sci 96:13203–13207.

Stan RV, Tkachenko E, Niesman IR. 2004. PV1 Is a Key Structural Component for the Formation of the Stomatal and Fenestral Diaphragms. Mol Biol Cell 15:3615–3630.

Tamoutounour S, Henri S, Lelouard H, Bovis B de, Haar C de, Woude CJ van der, Woltman AM, Reyal Y, Bonnet D, Sichien D, Bain CC, Mowat AM, Sousa CRe, Poulin LF, Malissen B, Guilliams M. 2012. CD64 distinguishes macrophages from dendritic cells in the gut and reveals the Th1-inducing role of mesenteric lymph node macrophages during colitis. Eur J Immunol [Internet] 42:3150–3166. Available from: http://doi.wiley.com/10.1002/eji.201242847

Trimm E, Red-Horse K. 2022. Vascular endothelial cell development and diversity. Nat Rev Cardiol:1–14.

Ubogu EE. 2020. Biology of the human blood-nerve barrier in health and disease. Exp Neurol 328:113272.

Vanlandewijck M, He L, Mäe MA, Andrae J, Ando K, Gaudio FD, Nahar K, Lebouvier T, Laviña B, Gouveia L, Sun Y, Raschperger E, Räsänen M, Zarb Y, Mochizuki N, Keller A, Lendahl U, Betsholtz C. 2018. A molecular atlas of cell types and zonation in the brain vasculature. Nature [Internet] 554:475–480. Available from: http://www.nature.com/articles/nature25739

Vlist M van der, Raoof R, Willemen HLDM, Prado J, Versteeg S, Gil CM, Vos M, Lokhorst RE, Pasterkamp RJ, Kojima T, Karasuyama H, Khoury-Hanold W, Meyaard L, Eijkelkamp N. 2022. Macrophages transfer mitochondria to sensory neurons to resolve inflammatory pain. Neuron 110:613–626.e9.

Wang PL, Yim AKY, Kim K-W, Avey D, Czepielewski RS, Colonna M, Milbrandt J, Randolph GJ. 2020a. Peripheral nerve resident macrophages share tissue-specific programming and features of activated microglia. Nat Commun 11:2552.

Wang Z, Liu C-H, Huang S, Fu Z, Tomita Y, Britton WR, Cho SS, Chen CT, Sun Y, Ma J, He X, Chen J. 2020b. Wnt signaling activates MFSD2A to suppress vascular endothelial transcytosis and maintain blood-retinal barrier. Sci Adv 6:eaba7457.

Willis CL, Garwood CJ, Ray DE. 2007. A size selective vascular barrier in the rat area postrema formed by perivascular macrophages and the extracellular matrix. Neuroscience 150:498–509.

Yang AC, Vest RT, Kern F, Lee DP, Agam M, Maat CA, Losada PM, Chen MB, Schaum N, Khoury N, Toland A, Calcuttawala K, Shin H, Pálovics R, Shin A, Wang EY, Luo J, Gate D, Schulz-Schaeffer WJ, Chu P, Siegenthaler JA, McNerney MW, Keller A, Wyss-Coray T. 2022. A human brain vascular atlas reveals diverse mediators of Alzheimer’s risk. Nature 603:885–892.

Ydens E, Amann L, Asselbergh B, Scott CL, Martens L, Sichien D, Mossad O, Blank T, Prijck SD, Low D, Masuda T, Saeys Y, Timmerman V, Stumm R, Ginhoux F, Prinz M, Janssens S, Guilliams M. 2020. Profiling peripheral nerve macrophages reveals two macrophage subsets with distinct localization, transcriptome and response to injury. Nat Neurosci 23:676–689.

Yim AKY, Wang PL, Bermingham JR, Hackett A, Strickland A, Miller TM, Ly C, Mitra RD, Milbrandt J. 2022. Disentangling glial diversity in peripheral nerves at single-nuclei resolution. Nat Neurosci 25:238–251.

Yona S, Kim K-W, Wolf Y, Mildner A, Varol D, Breker M, Strauss-Ayali D, Viukov S, Guilliams M, Misharin A, Hume DA, Perlman H, Malissen B, Zelzer E, Jung S. 2013. Fate Mapping Reveals Origins and Dynamics of Monocytes and Tissue Macrophages under Homeostasis. Immunity [Internet] 38:79–91. Available from: http://linkinghub.elsevier.com/retrieve/pii/S1074761312005481

Yu X, Liu H, Hamel KA, Morvan MG, Yu S, Leff J, Guan Z, Braz JM, Basbaum AI. 2020. Dorsal root ganglion macrophages contribute to both the initiation and persistence of neuropathic pain. Nat Commun [Internet] 11:264. Available from: http://www.nature.com/articles/s41467-019-13839-2

Zhang H, Li Y, Carvalho-Barbosa M de, Kavelaars A, Heijnen CJ, Albrecht PJ, Dougherty PM. 2016. Dorsal Root Ganglion Infiltration by Macrophages Contributes to∼Paclitaxel Chemotherapy-Induced Peripheral Neuropathy. J Pain [Internet] 17:775–786. Available from: http://linkinghub.elsevier.com/retrieve/pii/S1526590016005629

Zhang W, Dai M, Fridberger A, Hassan A, DeGagne J, Neng L, Zhang F, He W, Ren T, Trune D, Auer M, Shi X. 2012. Perivascular-resident macrophage-like melanocytes in the inner ear are essential for the integrity of the intrastrial fluid–blood barrier. Proc National Acad Sci 109:10388–10393.

Zhao Z, Nelson AR, Betsholtz C, Zlokovic BV. 2015. Establishment and Dysfunction of the Blood-Brain Barrier. Cell 163:1064–1078.

Zigmond RE, Echevarria FD. 2019. Macrophage biology in the peripheral nervous system after injury. Prog Neurobiol [Internet] 173:102–121. Available from: https://linkinghub.elsevier.com/retrieve/pii/S030100821830128X

