## Supplementary Figures for "A network of CD163^+^ macrophages monitors enhanced permeability at the blood-dorsal root ganglion barrier"

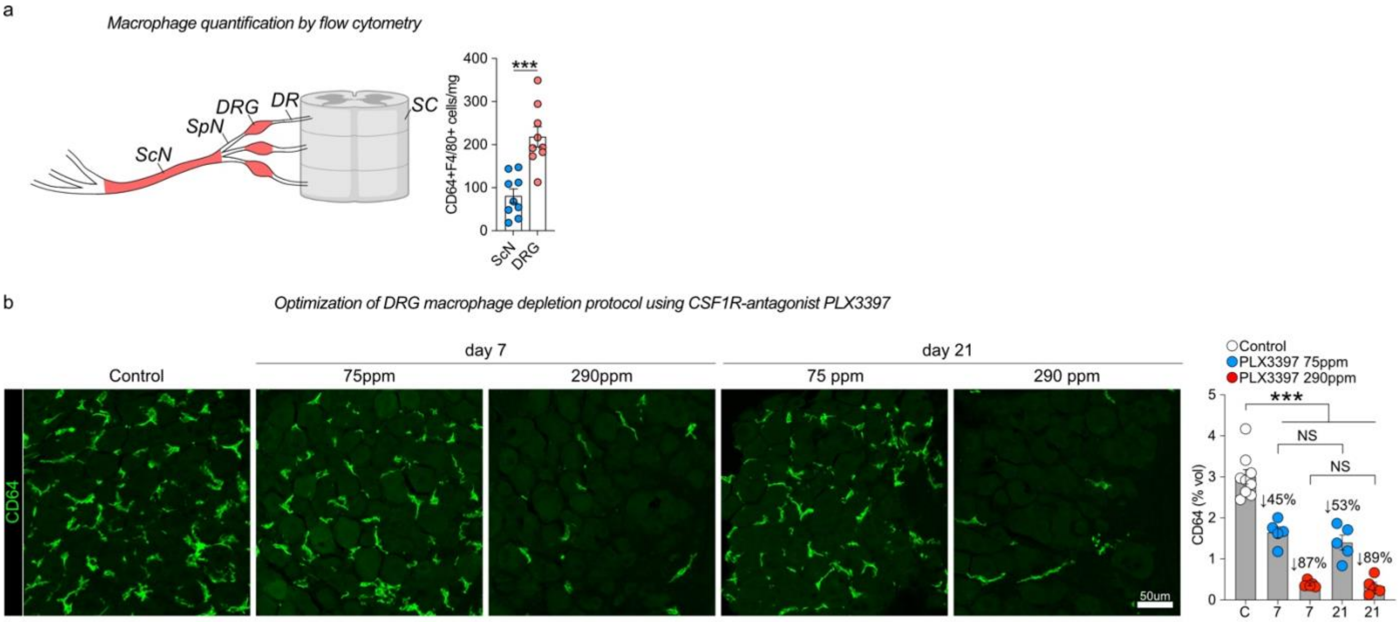


**Supplementary Figure 1 related to Figure 1**

(a) Flow cytometry based quantification of $CD64^{+}F4/80^{+}$ DRG or ScN macrophages, normalized to wet weight of tissue. n= 3 experiments pooled, 3mice/exp. Student's unpaired t-test.

(b) Quantification of $CD64^{+}$ DRG macrophages in mice administered chow containing the CSF1R antagonist PLX3397 at 75 ppm or 290 ppm for 7 or 21 days. 90% depletion is already achieved with the 290 ppm dose after 7 days. n=10, 5, 5 mice/group. The 75 ppm dose was performed once. The 290ppm dose was performed twice. Tukey's multiple comparisons test.

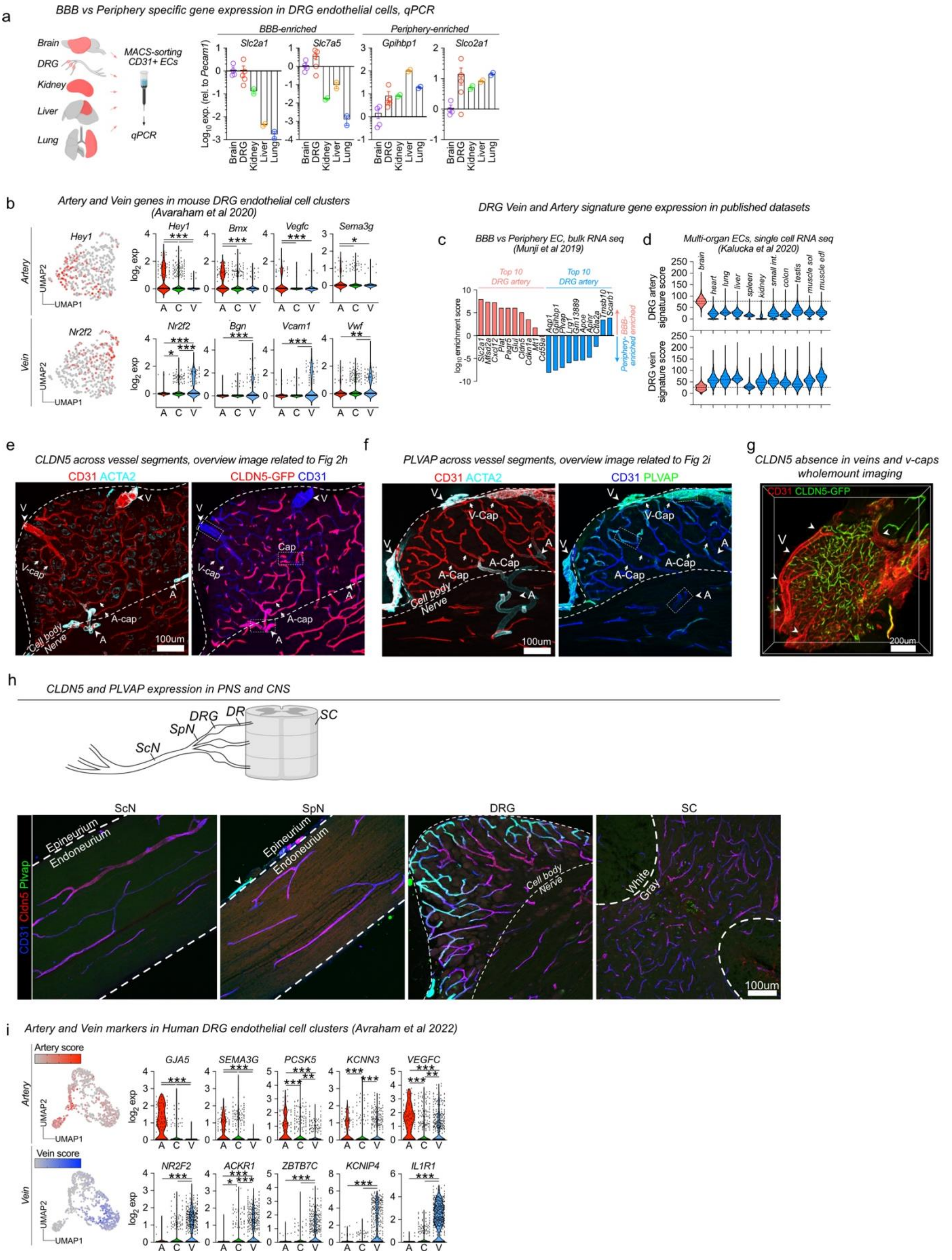


**Supplementary Figure 2 related to Figure 2**

(a) qPCR of MACS-enriched CD31$^{+}$ endothelial cells from indicated organs, normalized to brain.

(b) Expression of vein or artery signature genes across mouse DRG endothelial clusters (Avraham et al 2020).

(c) Enrichment of DRG vein or artery signature genes in BBB or peripheral endothelial cells (Munji et al 2019). DRG artery genes are enriched in BBB, whereas DRG vein genes are enriched in peripheral endothelium.

(d) Expression of DRG vein or artery signature genes in endothelial cells isolated from indicated organs (Kalucka et al 2020). DRG artery genes are enriched in brain endothelial cells, and DRG vein genes are most enriched in liver and muscle endothelium.
(e) CLDN5-GFP expression in DRG section, visualizing cropped areas in Fig 2h.
(f) Immunostaining of PLVAP in DRG section, visualizing cropped areas in Fig 2i.
(g) Wholemount imaging of native GFP expression in DRGs from *Cldn5*$^{gfp/+}$ mice, confirming absence of CLDN5 expression in large veins and venous capillaries.
(h) CLDN5 and PLVAP immunostaining in indicated tissues. Outside of the DRG, PLVAP$^{+}$CLDN5$^{-}$ vessels are only present in epineurial blood vessels. ScN and SpN endoneurial vessels are PLVAP$^{-}$CLDN5$^{+}$.
(i) Artery and Vein signature gene expression in human DRG endothelial clusters (Avraham et al 2022). Tukey's multiple comparisons test.

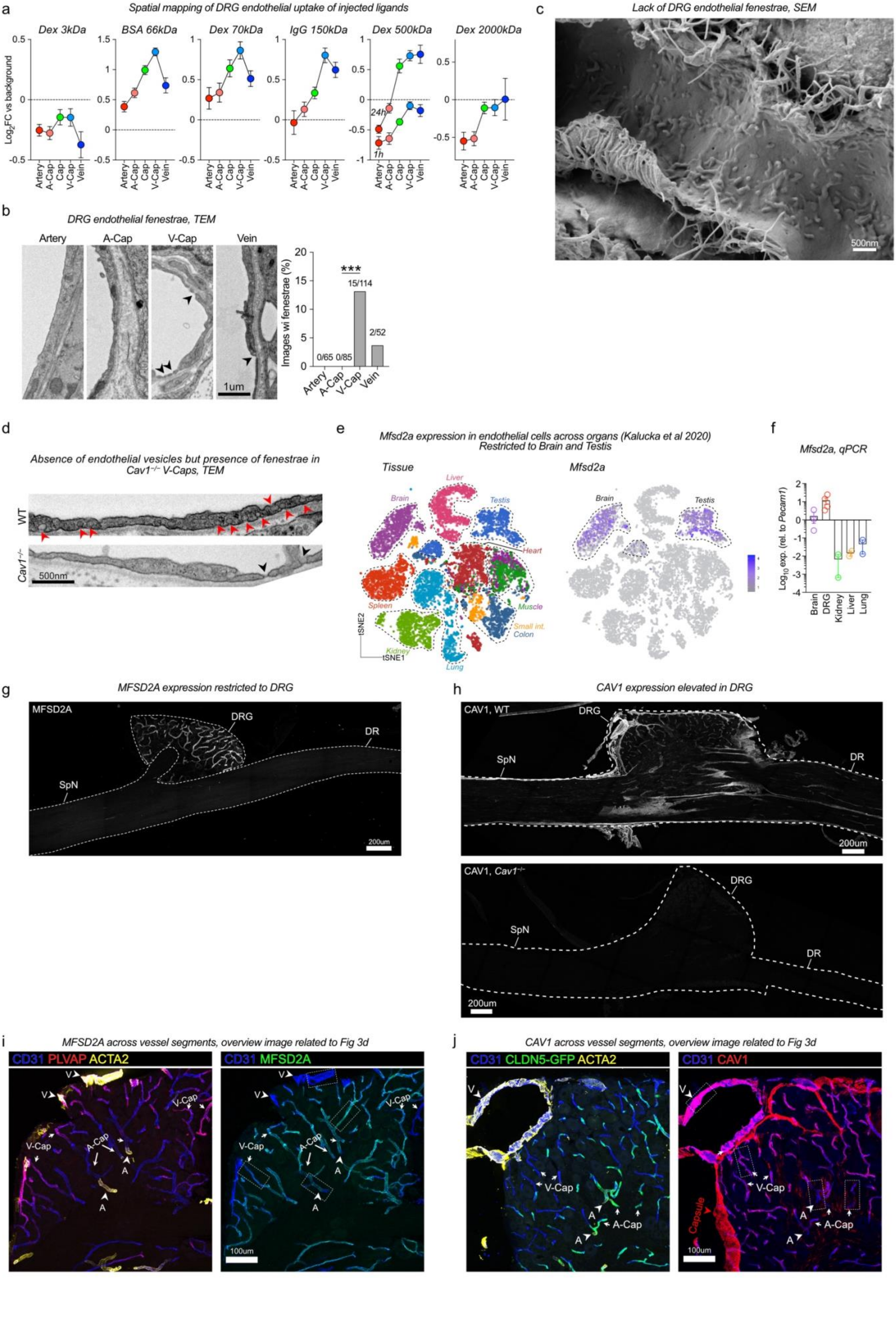
a
Spatial mapping of DRG endothelial uptake of injected ligands
Dex 3kDa
BSA 66kDa
Dex 70kDa
IgG 150kDa
Dex 500kDa
Dex 2000kDa
Log2FC vs background
24h
1h
Artery
A-Cap
Cap
V-Cap
Vein
b
DRG endothelial fenestrae, TEM
Artery
A-Cap
V-Cap
Vein
1um
Images wi fenestrae (%)
***
0/65
0/85
15/114
2/52
c
Lack of DRG endothelial fenestrae, SEM
500nm
d
Absence of endothelial vesicles but presence of fenestrae in Cav1−/− V-Caps, TEM
WT
Cav1−/−
500nm
e
Mfsd2a expression in endothelial cells across organs (Kalucka et al 2020)
Restricted to Brain and Testis
Tissue
Mfsd2a
Liver
Brain
Testis
Heart
Spleen
Muscle
Small int.
Colon
Kidney
Lung
tSNE1
tSNE2
f
Mfsd2a, qPCR
Log10 exp. (rel. to Pecam1)
Brain
DRG
Kidney
Liver
Lung
g
MFSD2A expression restricted to DRG
MFSD2A
DRG
DR
SpN
200um
h
CAV1 expression elevated in DRG
CAV1, WT
DRG
SpN
DR
200um
CAV1, Cav1−/−
DRG
SpN
DR
200um
i
MFSD2A across vessel segments, overview image related to Fig 3d
CD31 PLVAP ACTA2
CD31 MFSD2A
V
V-Cap
A-Cap
A
100um
j
CAV1 across vessel segments, overview image related to Fig 3d
CD31 CLDN5-GFP ACTA2
CD31 CAV1
V
V-Cap
A-Cap
A
Capsule
100um

**Supplementary Figure 3 related to Fig 3**

(a) Uptake of indicated i.v-injected tracers in DRG endothelial cells, separated by vessel segments. The following tracers, doses and circulation times were used3kDa dextran-TMR (25mg/kg, 1h, n=4 mice), BSA-A647 (5mg/kg, 1h, n=4 mice), 70kDa dextran TMR (25mg/kg, 1h, n=9 mice), goat anti rabbit IgG-A488 (4mg/kg, 4h, n=4 mice), 500kDa dextran-FITC (25mg/kg, 1h or 24h, n=4 mice), 2000kDa dextran-FITC (25mg/kg, 24h, n=4 mice). Values are mean of individual macrophages, normalized to tissue background.

(b) Analysis of endothelial fenestrae in TEM images from indicated DRG vessel segments. Total images per vessel segment indicated above each bar. n=4 mice. Fisher's exact test.

(c) Scanning electron micrograph of luminal surface of DRG v-cap with an apparent scarcity of endothelial fenestrae.

(d) Representative TEM images of v-caps from WT and *Cav1*$^{-/-}$ mice, showing the absence of caveolar vesicles (red arrowhead), but presence of fenestrae (black arrowhead) in *Cav1*$^{-/-}$ mice.

(e) *Mfsd2a* expression across endothelial cells isolated from multiple organs showing expression restricted to brain and testis.

(f) *Mfsd2a* expression in MACS-enriched CD31$^{+}$ endothelial cells from indicated organs, normalized to brain.

(g) MFSD2A immunostaining in DRG section.

(h) CAV1 immunostaining in entire DRG section from WT and *Cav1*$^{-/-}$ mice, demonstrating antibody specificity.

(i) Representative MFSD2A immunostaining in DRG section, visualizing cropped areas in Fig 3d.

(j) Representative CAV1 immunostaining in DRG section, visualizing cropped areas in Fig 3d.

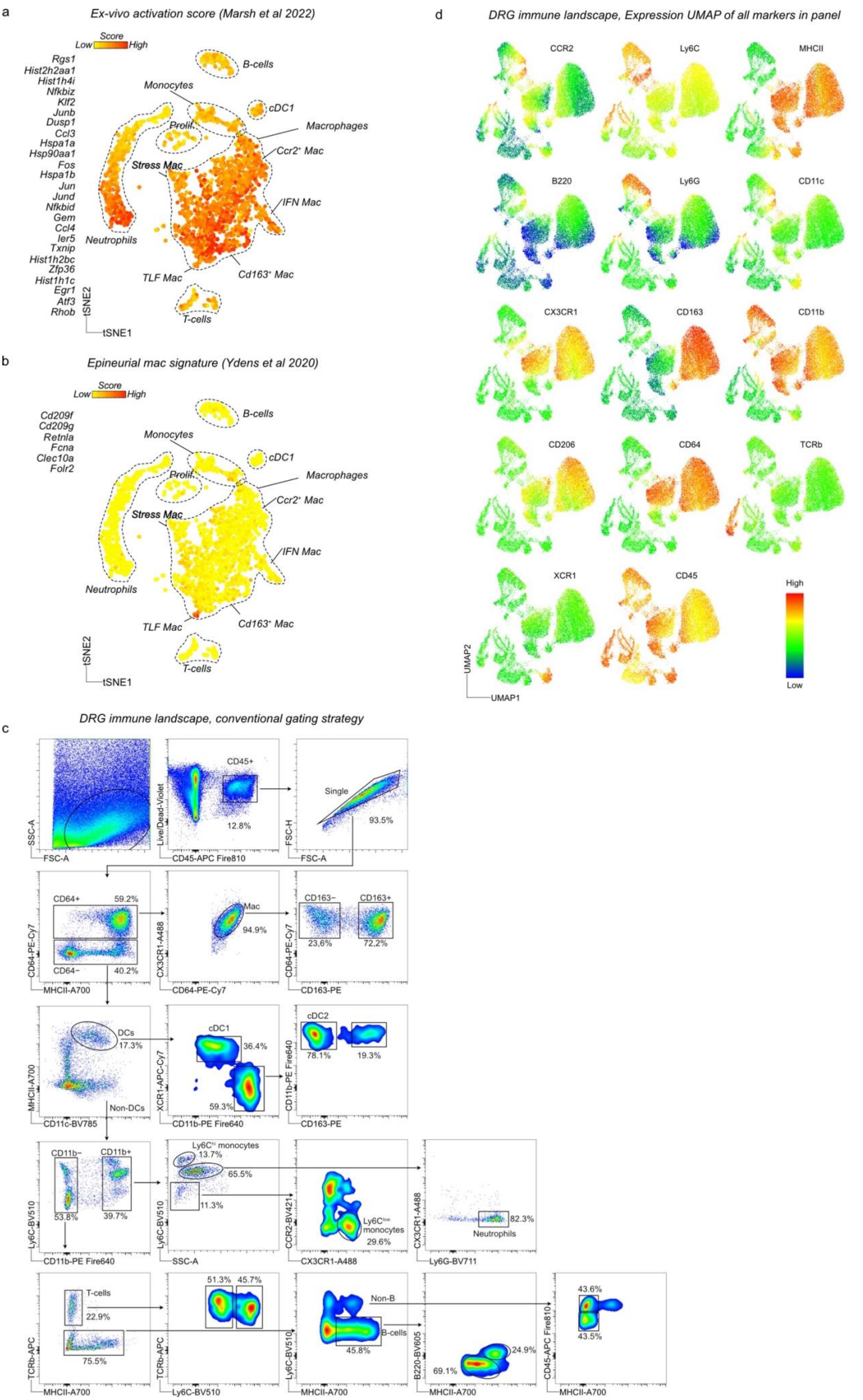

a
Ex-vivo activation score (Marsh et al 2022)
Score
Low
High
Rgs1
Hist2h2aa1
Hist1h4i
Nfkbiz
Klf2
Junb
Dusp1
Ccl3
Hspa1a
Hsp90aa1
Fos
Hspa1b
Jun
Jund
Nfkbid
Gem
Ccl4
Ier5
Txnip
Hist1h2bc
Zfp36
Hist1h1c
Egr1
Atf3
Rhob
B-cells
Monocytes
cDC1
Macrophages
Prolif.
Ccr2+ Mac
Stress Mac
IFN Mac
Neutrophils
TLF Mac
Cd163+ Mac
T-cells
tSNE2
tSNE1
b
Epineurial mac signature (Ydens et al 2020)
Score
Low
High
Cd209f
Cd209g
Retnla
Fcna
Clec10a
Folr2
B-cells
Monocytes
cDC1
Macrophages
Prolif.
Ccr2+ Mac
Stress Mac
IFN Mac
Neutrophils
TLF Mac
Cd163+ Mac
T-cells
tSNE2
tSNE1
c
DRG immune landscape, conventional gating strategy
SSC-A
FSC-A
CD45+
12.8%
Live/Dead-Violet
CD45-APC Fire810
Single
93.5%
FSC-H
FSC-A
CD64+
59.2%
CD64−
40.2%
CD64-PE-Cy7
MHCII-A700
Mac
94.9%
CX3CR1-A488
CD64-PE-Cy7
CD163−
CD163+
23,6%
72,2%
CD64-PE-Cy7
CD163-PE
DCs
17.3%
Non-DCs
MHCII-A700
CD11c-BV785
cDC1
36.4%
59.3%
XCR1-APC-Cy7
CD11b-PE Fire640
cDC2
78.1%
19.3%
CD11b-PE Fire640
CD163-PE
CD11b−
CD11b+
53.8%
39.7%
Ly6C-BV510
CD11b-PE Fire640
Ly6Chi monocytes
13.7%
65.5%
11.3%
Ly6C-BV510
SSC-A
Ly6Clow monocytes
29.6%
CCR2-BV421
CX3CR1-A488
82.3%
Neutrophils
CX3CR1-A488
Ly6G-BV711
T-cells
22.9%
75.5%
TCRb-APC
MHCII-A700
51.3%
45.7%
TCRb-APC
Ly6C-BV510
Non-B
B-cells
45.8%
Ly6C-BV510
MHCII-A700
24.9%
69.1%
B220-BV605
MHCII-A700
43.6%
43.5%
CD45-APC Fire810
MHCII-A700
d
DRG immune landscape, Expression UMAP of all markers in panel
CCR2
Ly6C
MHCII
B220
Ly6G
CD11c
CX3CR1
CD163
CD11b
CD206
CD64
TCRb
XCR1
CD45
High
Low
UMAP2
UMAP1

**Supplementary Figure 4 related to Fig 4**

(a) *Ex vivo* activation gene signature (25 genes identified in Marsh et al 2022) across DRG immune cell types. Induction of gene signature is most apparent in macrophages.
(b) Epineurial macrophage signature (selected genes from Ydens et al 2020) across DRG immune cell types. The signature is restricted to TLF macrophages.
(c) Traditional gating strategy to identify major immune cell populations in the UMAP.
(d) Expression heatmap of all markers across UMAP clusters.

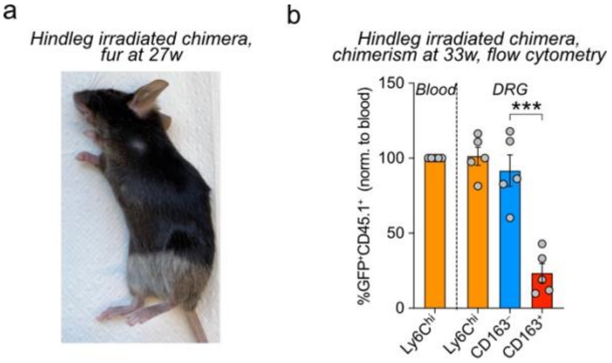


**Supplementary Figure 5 related to Fig 5**

(a) Mouse fur at 27 weeks after hindleg irradiation.
(b) Flow cytometry-based analysis of chimerism at 33 weeks after irradiation. Student's unpaired t-test.
